# Profiling the CFTR Variant Selectivity and Off-Target Interactions of VX-121

**DOI:** 10.64898/2026.06.01.729306

**Authors:** Ashish R. Jhangiani, John A. Olson, Austin Tedman, Catherine Foye, JaNise J. Jackson, Ashlyn G. Winters, Janiyah A. White, Mia Perfetti, Grace M. Abell, Crissey D. Cameron, Liudmyla Arifova, Brianna Corman, J. Paul Robinson, Charles P. Kuntz, Kaitlyn V. Ledwitch, Jens Meiler, Kathryn E. Oliver, Lars Plate, Jonathan P. Schlebach

**Author notes:** Authors contributed equally (Ashish R. Jhangiani, John A. Olson).

## Abstract

More than 1,200 variants of the cystic fibrosis transmembrane conductance regulator gene (*CFTR*) are associated with cystic fibrosis (CF), an autosomal recessive pulmonary disease affecting over 100,000 people. Most people with CF bear a common *CFTR* variant (F508del) that can be treated with therapeutics containing “correctors” that suppress the misfolding of the CFTR chloride channel. However, the pharmacological responsiveness of other rare CF variants can vary tremendously. The approval of VX-121, a VX-445 analog that serves as a key component of Alyftrek^TM^, potentially provides a new therapeutic option for those with rare CF variants. Nevertheless, it remains unclear whether VX-121 offers superior rescue across the entire spectrum of rare CF variants. In this work, we use deep mutational scanning (DMS) to survey the impact of VX-121 on the plasma membrane expression of 232 rare CF variants. Our results show that VX-121 generally enhances CF variant expression more than VX-445 and is most potent towards variants with mutations in the first membrane spanning domain (MSD1). However, we identify one variant (Y1032C) with diminished proteostatic and functional selectivity for VX-121 relative to VX-445. Computational docking suggests that the native Y1032 side chain forms favorable interactions with VX-121 that are disrupted by this mutation in a manner that alters its coordination. Finally, using photo-crosslinking, we show that VX-121 avoids a key off-target interaction of VX-445. Together, our findings provide new insights into the similarities and differences between current approved CF therapeutics.

## Introduction

Cystic fibrosis (CF) is an autosomal recessive disorder caused by loss-of-function variants in the CF transmembrane conductance regulator gene (*CFTR*), which encodes a chloride ion channel. This channel maintains the electrochemical potential of chloride and bicarbonate ions across secretory epithelial membranes within numerous tissues. Improper CFTR function results in physiological salt imbalances that ultimately cause a variety of clinical issues, such as exocrine pancreatic insufficiency, infertility, and chronic susceptibility to pulmonary infections. CF most often arises from the effects of one or more copies of the c.1521_1523delCTT variant (p.PheF508del or F508del), which continues to persist with high penetrance. Nevertheless, thousands of more rare CF variants have been identified to date (wwww.cftr2.org/). (1) While this spectrum of loss-of-function variants causes similar clinical outcomes, the underlying molecular pathologies can vary in a manner that impacts how different people with CF respond to approved therapeutics.(2) Most CF variants exhibit enhanced misfolding and degradation of the immature CFTR protein in the endoplasmic reticulum but also have secondary functional defects at the plasma membrane.(2) Efforts to understand the molecular basis of drug-variant interactions and to optimize the matching of therapeutics to *CFTR* genotypes therefore address a key challenge in precision medicine for people with CF.(3)

Several highly efficacious small-molecule therapies have been approved for the treatment of CF and have had a remarkable clinical impact over the last decade, as tens of thousands of people with CF now live with radically improved clinical prognoses.(4) Leading therapeutics such as Orkambi^TM^, Symdeko^TM^, and Trikafta^TM^combine a small molecule “potentiator” (VX-770 or VX-561), which enhances CFTR gating, with one or more “corrector” molecules that suppress CFTR misfolding (e.g. VX-809 or VX-661, and VX-445 or VX-121).(5) Though these therapies generally deliver excellent efficacy for people bearing one or two F508del *CFTR* alleles,(6–8) they are not universally efficacious for those with other genotypes due to their inconsistent rescue of rare CF variants.(3, 9) We previously compared the effects of VX-661 and VX-445 on the plasma membrane expression of 232 clinical CFTR variants.(10, 11) While both correctors rescue the expression of a variety of CF variants, we found that the distinct binding modes gave rise to variant-specific differences in the magnitude of their rescue.(10) The United States Food and Drug Administration (FDA) has since approved Alyftrek^TM^,(7, 8) a combination therapy akin to Trikafta^TM^ that exchanges VX-445 for VX-121 and VX-770 for VX-561 (deuterated VX-770). While this therapy was generally well-tolerated and improved CFTR function among patients bearing at least one F508del variant,(7, 8) it remains unclear whether VX-121 exerts equivalent effects to VX-445 across the entire spectrum of rare CF variants.

In the following, we survey the effects of VX-121 and VX-445 on the expression and function of rare CF variants, then explore differences in their on- and off-target binding. We first employ deep mutational scanning (DMS) to measure the impact of VX-121 on the plasma membrane expression of 232 clinical missense and indel variants. Overall, we find the magnitude of the proteostatic correction generated by VX-121 to be greater than that of VX-445 for most rare CF variants. Like VX-445, our results reveal that VX-121 enhances the expression of a wide variety of variants and exhibits the highest potency towards variants with mutations in the first membrane spanning domain (MSD1) of CFTR. Nevertheless, our results identify one variant, c.3095A>G (p.Tyr1032Cys or Y1032C), with diminished selectivity for VX-121 and another that exhibits elevated rescue of expression in the presence of VX-121 (V1240G). Molecular docking investigations suggest VX-121 forms critical interactions with residue Y1032, and that the mutation of this residue alters the VX-121 binding mode. Interestingly, we find that Y1032C CFTR also exhibits functional selectivity for VX-445. Finally, using photo-crosslinking, we show that VX-121 is unlikely to form the same off-target interactions as VX-445. Together, our findings provide new insights into how modifications to the structure of correctors may alter variant-specific effects.

## Results

### Impact of VX-121 on the plasma membrane expression of CF Variants

We previously identified several instances in which certain CFTR variants preferentially respond to one corrector over another.(10) To survey the variant-specific effects of VX-121, we employed our previously described DMS platform to survey its effects on the plasma membrane expression of 232 CF variants.(10, 11) Briefly, we first generated a pool of recombinant HEK293T cells in which each cell expresses a single CF variant from a uniform genomic locus. We then employed CFTR surface immunostaining in conjunction with fluorescent cell sorting to separate the pool of cells according to the relative abundance of the CF variants expressed within each cell. Similar to our observations for VX-661 and VX-445,(10) the pool of recombinant cells expressing individual CF variants skews to higher surface CFTR immunostaining in the presence of VX-121 (Fig. 1A). Interestingly, VX-121 generates a larger increase in the mean fluorescence intensity (+112 ± 13%) relative to VX-445 (+51 ± 6%) under equivalent experimental conditions (Fig. 1A), which implies VX-121 is generally more potent towards rare variants overall. To compare the relative plasma membrane expression (PME) of each variant in the presence or absence of VX-121, we used deep sequencing to track the relative abundance of each variant within each cellular isolate and to estimate their corresponding surface immunostaining intensities. Variant immunostaining intensities upon addition of VX-121 are highly correlated with those from previous DMS measurements recorded in the presence of VX-445 (Spearman’s ρ = 0.96, *p* = 4.5*10^-174^) and to a lesser extent VX-661 (Spearman’s ρ = 0.90, *p* = 7.0*10^-120^, Fig. S1). These observations demonstrate that VX-121 corrects a similar sub-set of rare variants that are sensitive to other correctors.

**Figure 1.**
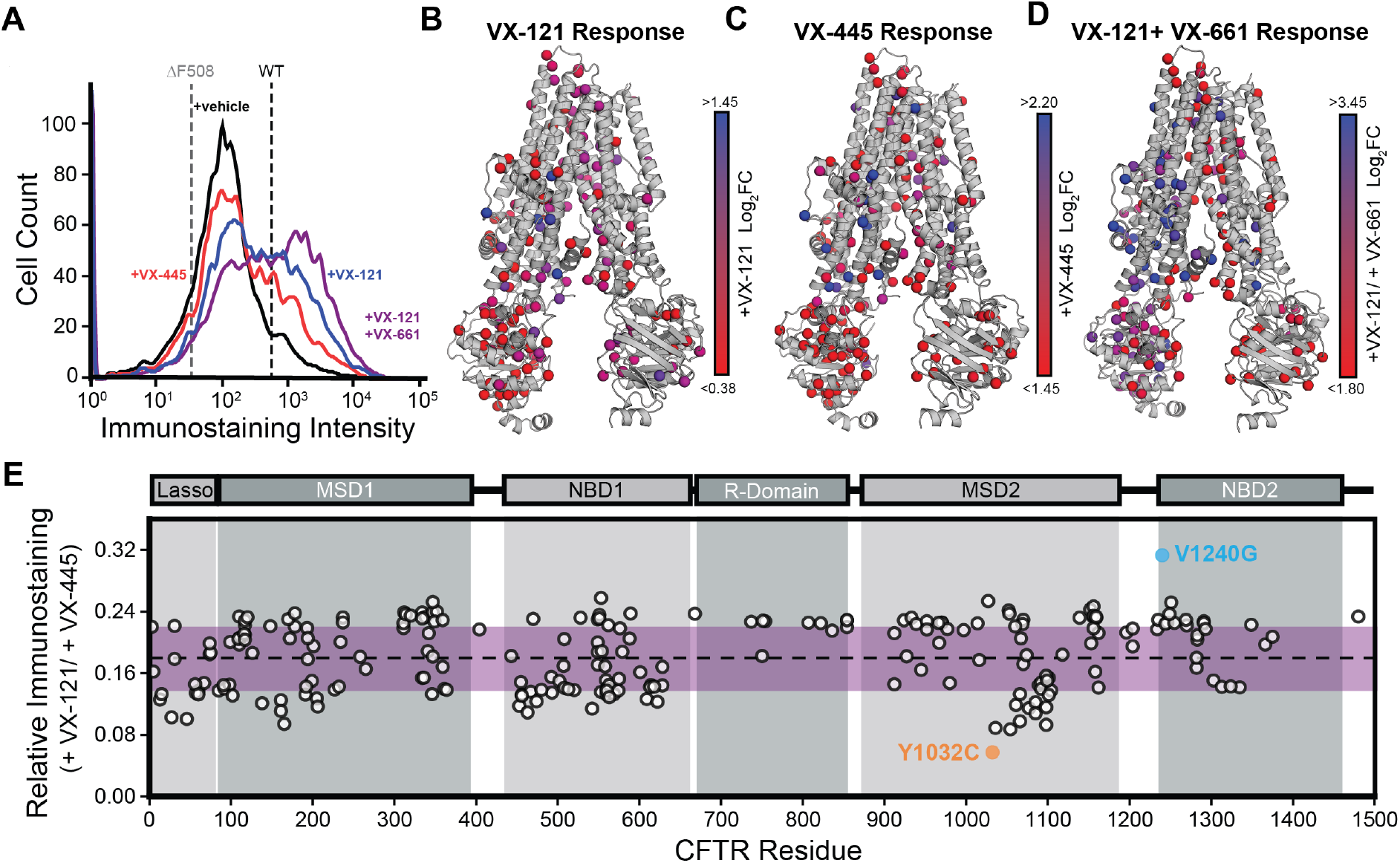
Impact of VX-121 on the Plasma Membrane Expression of CF Variants. Deep mutational scanning was employed to compare the impacts of VX-121 and various other correctors on the plasma membrane expression of 232 rare CF variants. A) A histogram depicts the distribution of CFTR surface immunostaining intensities among recombinant cells expressing individual CF variants in the presence of vehicle (black), 3 µM VX-121 (blue), 3 µM VX-445 (red) or 3 µM VX-121 + 3 µM VX-661 (purple). The approximate positions of the mean surface immunostaining intensities of recombinant cells expressing WT (black) or F508del CFTR (gray) are shown for reference. The log_2_FC in surface immunostaining of each variant in the presence of B) 3 µM VX-121, C) 3 µM VX-445, or D) 3 µM VX-121 and 3 μM VX-661 are projected onto their corresponding residues within a structural model of the inactive conformation of CFTR (PDB 5UAK). E) CFTR surface immunostaining intensities for each variant in the presence of vehicle were used to scale the corresponding intensity values in the presence of 3 µM VX-121 or 3 µM VX-445 (see *Materials and Methods*). The ratio of the VX-121-treated intensity to the corresponding VX-445-treated intensity for each variant is plotted against the position of the mutated residues within the CFTR sequence. The boundaries of the lasso motif, membrane spanning domain 1 (MSD1), nucleotide binding domain 1 (NBD1), the R-domain, membrane spanning domain 2 (MSD2), and nucleotide binding domain 2 (NBD2) are highlighted in gray for reference, and the purple stripe reflects values within one standard deviation of the mean.

As is true for VX-445,(10) the fold-change in the immunostaining of individual variants suggests VX-121 sensitivity is maximized among correctable MSD1 variants near the VX-445 binding pocket (Table S1, Fig. 1 B-C), though allosteric correction across domains is achieved when combined with VX-661 (Fig. 1D). Direct comparison of DMS measurements with VX-121 to previously reported DMS measurements collected in the presence of VX-445 is complicated by differences in the dynamic range and fluorescence intensity scales associated with each DMS experiment. Therefore, to compare trends in corrector response, we scaled the VX-121-treated immunostaining intensity values for each variant according to the ratio of its vehicle-treated intensity values within each data set (see *Materials and Methods*). Dividing these scaled intensities in the presence of VX-121 by the corresponding values in the presence of VX-445 reveals which variants exhibit deviations in their response to one compound relative to the other (Fig. 1E). While the ratios for most variants fall within one standard deviation of the mean, there is a cluster of variants within MSD2 that appear to exhibit lower selectivity towards VX-121 (Fig. 1E). Notably, these variants bear mutations to residues within transmembrane domains (TMDs) 10 and 11, which form domain-swapped contacts within MSD1 and form the VX-445 binding pocket.(12– 15) The Y1032C variant appears to exhibit the weakest overall selectivity towards VX-121. In contrast, the V1240G variant appears to be slightly more sensitive to VX-121 relative to VX-445 (Fig. 1E). Though this side chain is distant from the VX-445 binding site (Fig. 2A), its apparent sensitivity to these correctors echoes our previous structural investigations suggesting VX-445 binding stabilizes the native conformation of nucleotide binding domain 2 (NBD2).(11) Together, these subtle differences potentially suggest differences in how they engage the CFTR protein.

**Figure 2.**
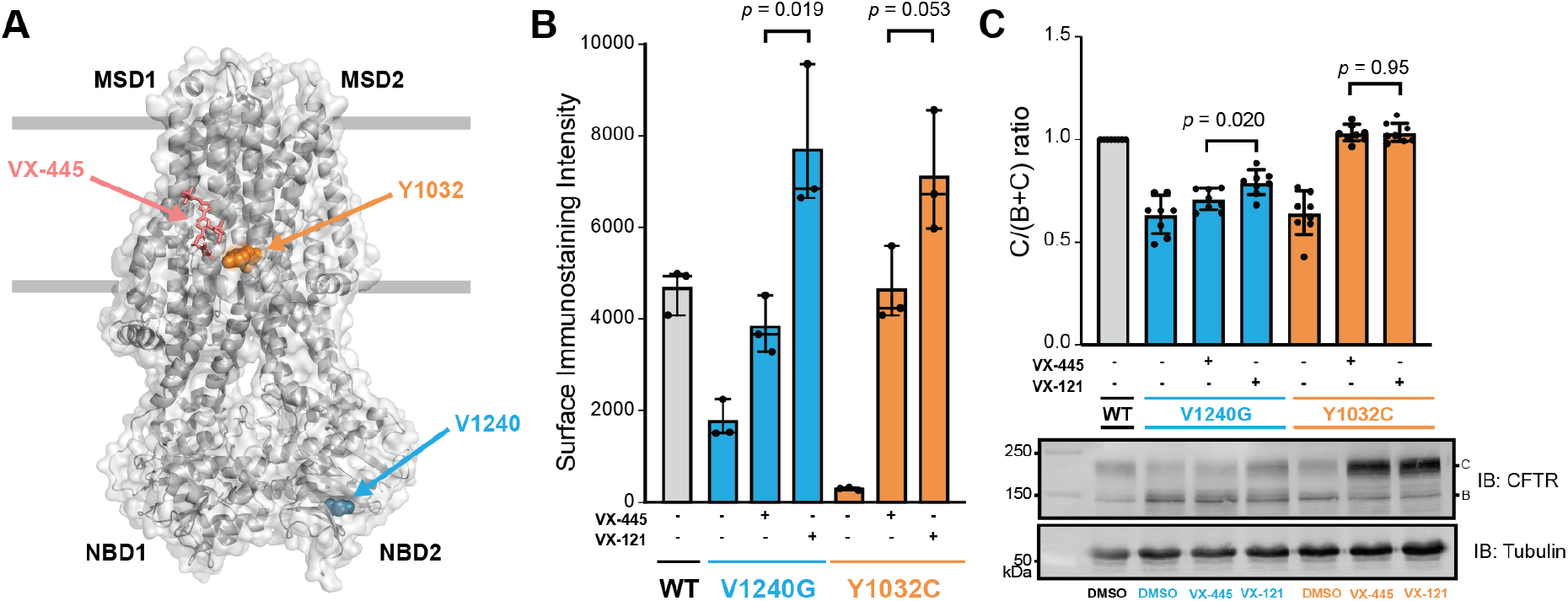
Proteostatic Response of V1240G and Y1032C CFTR to Correctors. Surface immunostaining and western blotting were used to quantitatively assess the effects of VX-121 and VX-445 on the plasma membrane expression and maturation of V1240G and Y1032C CFTR, respectively. A) A structural model of CFTR depicts the positions of the native Y1032 (orange) and V1240 (blue) side chains in relation to the position of the bound VX-445 (pink). B) Flow cytometry was used to quantitatively compare the relative surface immunostaining intensity of WT (gray), V1240G (blue), or Y1032 (orange) CFTR in stable HEK293T cell lines treated with vehicle, 3 µM VX-121, or 3 µM VX-445. A bar graph depicts the average surface immunostaining intensities across three biological replicates under each condition. Error bars reflect the standard deviations, and p-values from t-tests are shown. C) Western blots were used to quantitatively compare the relative maturation efficiency of transiently expressed WT (gray), V1240G (blue), or Y1032C (orange) CFTR in HEK293T cells treated with vehicle, 3 µM VX-121, or 3 µM VX-445. A representative blot shows the relative positions of mature (band C) and immature (band B) CFTR, and a tubulin loading control is shown for reference. A bar graph depicts the average ratio of the C band to total CFTR across eight biological replicates under each condition. Error bars reflect the standard deviations, and *p*-values from t-tests are shown.

### VX-121-mediated Correction of Y1032C and V1240G CFTR

While DMS measurements capture precise trends in variant surface immunostaining, the underlying raw intensity values determined with this approach are generally semi-quantitative.(10) Therefore, to validate the observed deviations in corrector selectivity, we quantified the effects of VX-121 and VX-445 on the surface immunostaining and cellular maturation of Y1032C and V1240G CFTR (Fig. 2A). We first used flow cytometry to quantify the surface immunostaining intensity of stable cells expressing each variant in the presence and absence of each corrector. Consistent with the wider overall trends for the surface immunostaining of recombinant cellular libraries (Fig. 1A), we find that the surface immunostaining of both variants is higher in the presence of VX-121 relative to VX-445 (Fig. 2B). By comparison, the difference in the immunostaining intensity of Y1032C CFTR treated with these two compounds is smaller (*p* = 0.053, Fig. 2B) than for V1240G (*p* = 0.019, Fig. 2B), which is consistent with the observed trends in selectivity. To compare how these compounds impact variant maturation, we used western blotting to measure the impact of each drug on the relative abundance of mature and immature CFTR levels. Though differences are subtle, a quantitative comparison of the ratio of the mature (C) and immature (B) glycoforms confirms that VX-121 generates more efficient rescue than VX-445 for V1240G CFTR, but not for Y1032C CFTR (Fig. 2C). Given that these two compounds bind within the same pocket with comparable affinity, these differences potentially reflect variations in the mode of corrector binding and stabilization.

### Structural basis of CF variant rescue by VX-121

Given that residue 1240 is distant from the VX-445 binding pocket, variations in the response of V1240G CFTR to these compounds likely involve differences in the efficiency of “allosteric” correction.(16) In contrast, previous cryo-EM structures show that the Y1032 side chain directly participates in the coordination of VX-445 within its binding pocket.(15) Given that VX-121 is a structural analog of VX-445, these apparent variations in the response of Y1032C CFTR could potentially reflect differences in how these molecules are coordinated by the CFTR protein. To investigate the molecular interactions governing VX-121 binding, we first used RosettaLigand and an open-state, VX-445-bound CFTR model to generate an ensemble of VX-121 binding poses. Docked poses were ranked by interface energy, and a representative high-confidence binding mode was identified through per-residue energy decomposition and manual inspection of the top-scoring models (Fig. 3A). Notably, the binding pocket for our top docked VX-121 poses is quite similar to the pocket of the VX-445-bound structure (Cα RMSD ∼0.7 Å), which is reasonable given that VX-121 is a VX-445 analog. Our top-selected model reveals a complex interaction network centered within the transmembrane pocket and lasso motif (interface energy = −16.8 Rosetta Energy Units, REU). In this pose, Y1032 emerged as the primary energetic contributor to the interface, providing a highly favorable per-residue interaction energy of −2.39 REU. While other high-ranking poses exhibited steric clashes or unfavorable energetics at this position, this model was prioritized for its engagement between the Y1032 side chain and the VX-121 scaffold. Analysis of the binding interface identified a network of interface residues that includes R21, R25, R29, and R1102, which together coordinate the ligand within the pocket (Fig. 3A). This predicted binding mode is consistent with our DMS data-the observed loss of VX-121 responsiveness in the Y1032C variant supports the computational finding that Y1032 is a critical anchor for VX-121 binding.

**Figure 3.**
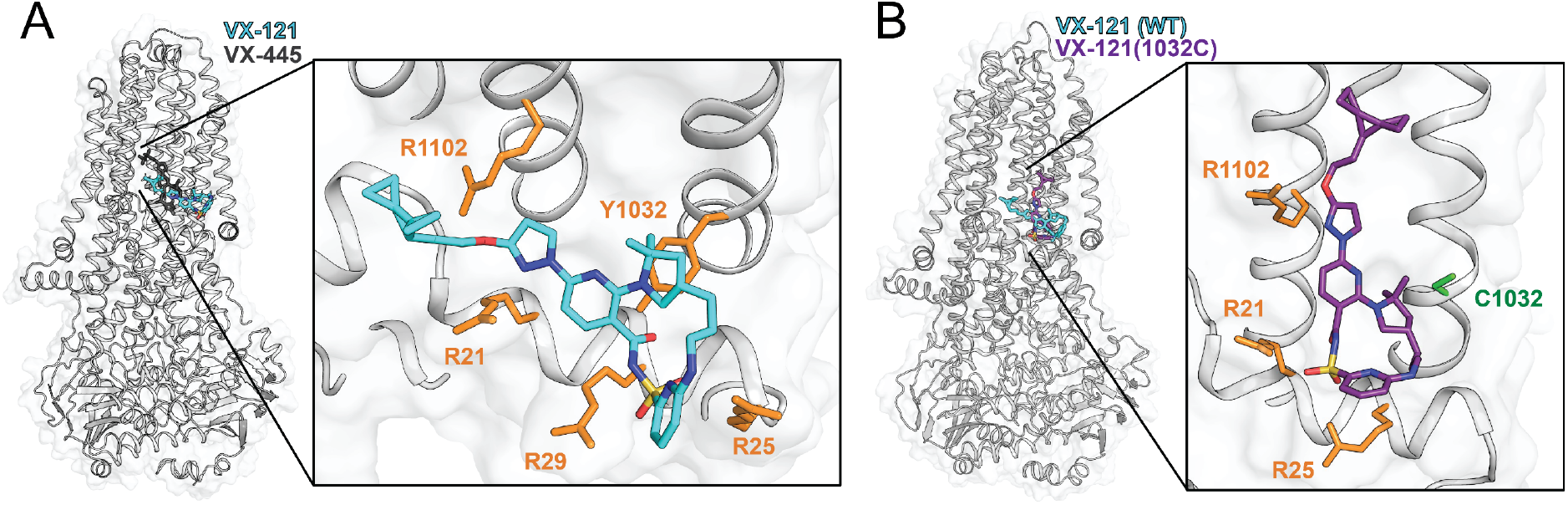
The Y1032 Side Chain Serves as a Key Structural Anchor for VX-121. Computational modeling was used to compare the manner in which CFTR coordinates VX-445 and VX-121, as well as how this is altered by mutations. A) The top-scoring docked pose for VX-121 (*cyan*) within the VX-445 binding pocket of WT CFTR (PDB 8EIG) is shown for reference. The experimental coordinates of the analog VX-445 (dark gray) are overlaid for reference. The inset highlights the predicted interaction network between VX-121 and key coordinating residues, including the Y1032 aromatic anchor and a cluster of arginine residues (orange). B) Comparative docking was used to assess how VX-121 is coordinated in the context of a structural model of Y1032C CFTR (purple). The top-scoring docked pose for VX-121 in the context of the WT CFTR model is shown for reference (cyan). The inset illustrates the change in the interaction network and the shift in ligand orientation upon substitution of the tyrosine anchor with cysteine (green). Key interface residues are shown in orange.

To assess the potential impact of the Y1032C mutant on this binding mode, we docked VX-121 against a model of the Y1032C variant and compared the top-scoring poses to wild-type (WT). The top mutant pose still engages R21, R25, and R1102, but lacks the native interaction with R29 (interface energy = −16.6 REU, Fig. 3B). Interestingly, the mutated C1032 forms an interaction with VX-121, though it is incapable of recapitulating the π stacking interaction that forms between VX-121 and Y1032 (Fig. 3). Per-residue energy decomposition across the top five mutant models showed that C1032 makes a smaller contribution to the interface energy (average −1.60 REU) across all five top poses relative to the native Y1032 side chain in the WT model (−2.39 REU). R25 and R21 were consistently favorable in the mutant (average −0.88 and −0.44 REU, respectively), whereas R1102 was strongly unfavorable (average +9.06 REU), which arises from a steric clash. Overall, the top mutant pose retains an interaction at position 1032 with the cysteine substitution, but with reduced energetics compared to WT. Whereas most variants exhibit a stronger response to VX-121 than VX-445, these observations suggest the Y1032C mutation perturbs VX-121 interactions in a manner that leads to a loss of this selectivity.

### Functional Correction of Y1032C and V1240G CFTR

Functional rescue of CF variants typically requires a combination of small molecules that restore folding and gating of the mature CFTR ion channel.(2) While both VX-121 and VX-445 can significantly enhance the plasma membrane expression of Y1032C and V1240G CFTR, it is unclear whether the observed correction results in recovery of CFTR activity. To gain insights into how these small molecules impact variant function, we measured short-circuit currents from Fischer rat thyroid (FRT) cell monolayers transiently expressing either Y1032C or V1240G CFTR. Impaired transepithelial ion transport is observed at baseline for both Y1032C (21.5 ± 1.3%) and V1240G (7.7 ± 1.4%) compared to wild-type (WT), which confirms these variants exhibit a loss-of-function. Nevertheless, the amplitude of the current generated by Y1032C CFTR indicates that it retains residual activity at the plasma membrane (Fig. 4A). Indeed, strong forskolin-stimulated rescue of Y1032C is observed in response to chronic treatment with any corrector (VX-661, VX-445, or VX-121) in the absence of a secondary potentiator (Fig. 4A). Interestingly, Y1032C currents are higher following treatment with VX-445 (91.2 ± 3.9% of WT) compared to VX-121 (57.3 ± 3.4% of WT), the latter of which generates similar functional rescue to the type I corrector VX-661 (52.4 ± 2.2% of WT) (Fig. 4A). This result implies VX-121 and VX-661 binding are incapable of stabilizing the native functional ensemble of Y1032C CFTR to the same extent as VX-445. Furthermore, structural perturbations of the binding pocket caused by the Y1032C variant result in greater correction and functional selectivity for VX-445 relative to VX-121.

**Figure 4.**
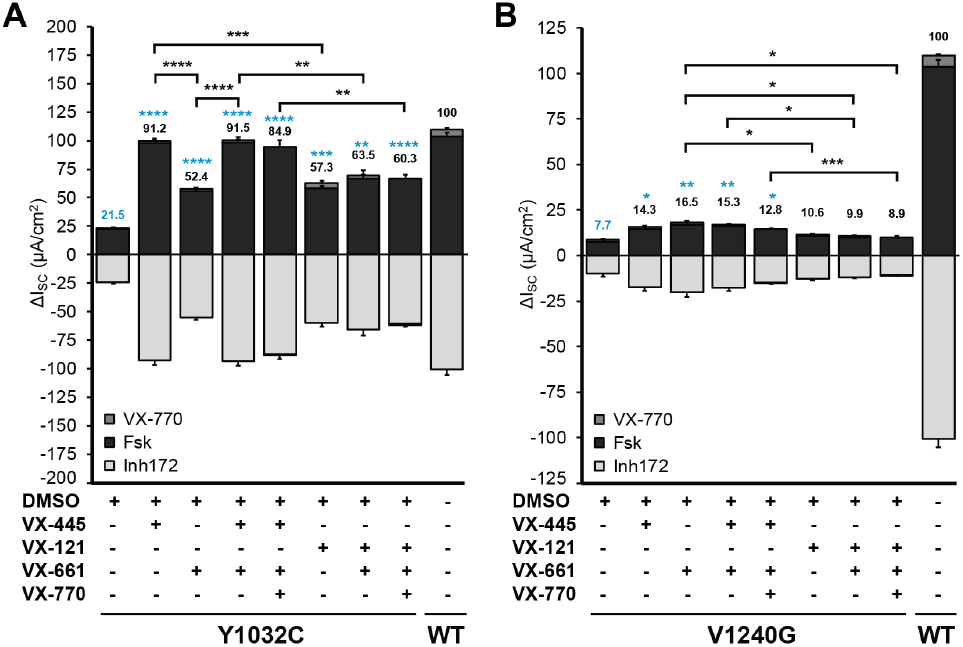
Functional Modulation of the Y1032C and V1240G Variants by CFTR Correctors and Potentiators. Ussing chamber results compare short-circuit currents across FRT monolayers transiently expressing (A) Y1032C or (B) V1240G. Cells are cultured for 5 days on permeable transwells and chronically treated with CFTR modulators (48hr, 5 µM per compound) or DMSO vehicle. Short-circuit currents (ΔI_SC_) are quantified following acute application of forskolin (Fsk; 10 µM), VX-770 (5 µM), and CFTR-specific Inhibitor-172 (Inh172, 10 µM). Deuterated VX-770 iwas used for treatment combinations with VX-121 and VX-661 to mimic the composition of Alyftrek^TM^. Data are mean ± SEM (n = 4-6 transwells per condition from 3 independent experiments). Numbers above each bar graph show average acute Fsk+VX-770 responses as percent of WT. Statistical comparisons are calculated based on Fsk+VX-770 values. *, p < 0.05; **, p < 0.01; ***, p < 0.001; ****, p < 0.0001 (one-way ANOVA). Blue asterisks indicate statistical comparison to DMSO.

Patterns of functional correction are distinctly different for V1240G CFTR, even though this CF variant exhibits comparable plasma membrane expression to Y1032C CFTR in the presence of either VX-121 or VX-445 (Fig. 2B). By comparison, V1240G CFTR exhibits much weaker functional rescue upon chronic treatment with single correctors, combinations of correctors, or combinations of correctors and potentiators. Like Y1032C CFTR, cocktails containing VX-445 generate significantly higher forskolin-stimulated currents (12.8 ± 0.6% to 15.3 ± 1.5% of WT) relative to those containing VX-121 (8.9 ± 0.5% to 10.6 ± 0.5% of WT) (Fig. 4B). Also of note, VX-661 alone augments V1240G activity to 16.5 ± 2.0% of WT, which is greater than VX-121 applied as a single agent (10.6 ± 0.5% of WT) or in combination with VX-661 (9.9 ± 0.5% of WT). Overall, findings indicate the attenuated proteostatic response of Y1032C to VX-121 translates to sizable functional selectivity for VX-445, while only VX-445 or VX-661 are able to generate meaningful functional rescue of V1240G CFTR.

### Off-target Interactions of VX-121

Trikafta^TM^ is associated with several known side effects and adverse events,(17–20) and multiple drug-drug interactions have been reported.(21–24) We recently found that VX-445 forms an off-target interaction with saccharopine dehydrogenase-like oxidoreductase (SCCPDH),(25) an enzyme that is possibly involved in neurotransmitter regulation. Though the side effect profiles of Trikafta^TM^ and Alyftrek^TM^ are broadly similar, it is unclear whether the structural differences between VX-121 and VX-445 could lead to distinct off-target interactions. To assess whether VX-121 also interacts with SCCPDH, we employed a functionalized photoaffinity ligand (PAL) of VX-445 (VU439, Fig. 5A). VU439 was added to HEK293T cells transiently expressing a Myc-tagged version of SCCPDH. The PAL was then cross-linked to interacting proteins prior to lysis and click chemistry-mediated functionalization of labeled proteins with the fluorescent handle TAMRA-azide-desthiobiotin. Fluorescently labeled, biotinylated proteins were then enriched using a streptavidin pulldown followed by visualization on a western blot (Fig 5B, left). Labeled lysates contain a prominent VU439-TMR-labeled SCCPDH band near 47 kDa, which was confirmed using an anti-Myc immunoblot (Fig. 5B, right). To compare the propensity of VX-445 and VX-121 to interact with this off-target, we repeated these experiments in the presence of competing unlabeled corrector molecules. The addition of excess unlabeled VX-445 decreased SCCPDH labeling by VU439 to levels that were undetectable by western blotting (Fig. 5B & C). In contrast, SCCPDH labeling was unaffected by the addition of excess VX-121, which suggests that VX-121 is incapable of outcompeting the probe. Together, these results highlight that VX-121 lacks a key off-target VX-445 interaction.

**Figure 5.**
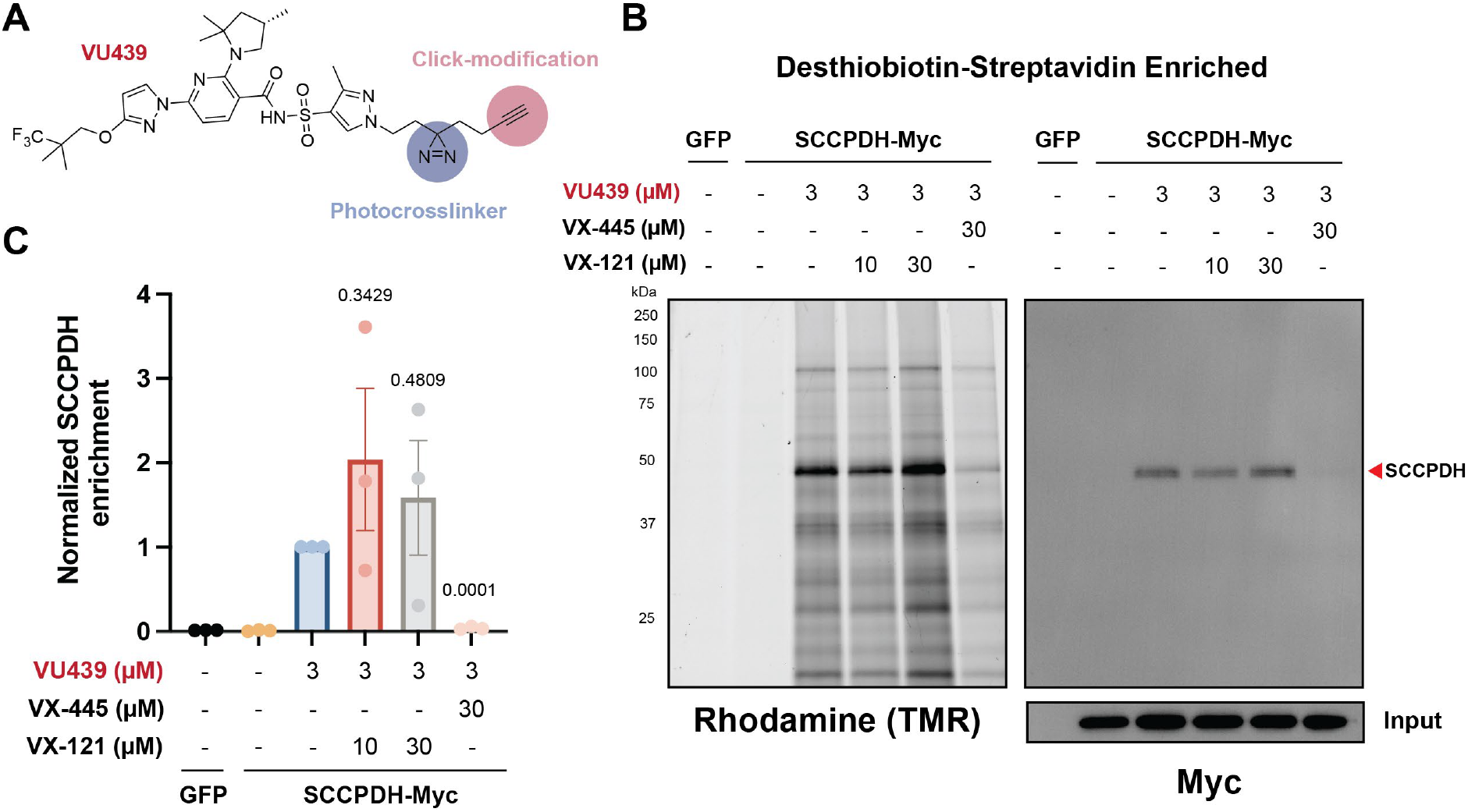
Off-Target Profiling of VX-121. A VX-445-based photo crosslinking probe (VU439) was used to compare the propensity of VX-121 and VX-445 to bind to the SCCPDH protein. A) HEK293T cells transiently expressing SCCPDH-Myc were treated with 3 µM VU439 probe prior to UV cross-linking, functionalization with TAMRA-azide-desthiobiotin, immunoprecipitation, and analysis via western blot. Fluorescent imaging of a representative blot shows the relative enrichment of lableled proteins (left) and an anti-Myc immunostaining of the blot confirms the identity of the major band of SCCPDH. B) The molecular structure of the photoaffinity ligand VU439 is shown for reference. The photocrosslinking and alkyne moiety used for click modifications are indicated. C) Photo crosslinking by VU439 was carried out in the presence of varying concentrations of unlabeled VX-445 or VX-121. A bar graph depicts the average enrichment of the SCCPDH protein relative to the VU439-only control across three biological replicates. Error bars reflect the standard deviation, and the p-values associated with a one-sample paired t-test are shown.

## Discussion

There are currently five FDA-approved CF drugs that can be used to treat hundreds of different *CFTR* genotypes, along with a variety of other small-molecule and nucleotide-based interventions that are undergoing clinical trials. While most people with CF derive immense clinical benefit from various combinations of CFTR modulators, many are left without viable therapeutic options for a multitude of reasons, including drug intolerance and/ or insufficient CFTR theratype response.(3) The recent approval of Alyftrek^TM^,(7, 8) which contains the VX-445 analog VX-121, provides an alternative therapeutic option for some individuals. Nevertheless, it remains unclear whether this treatment is broadly more efficacious across the spectrum of known CF variants. In this work, we employ DMS to compare the effects of VX-121 and VX-445 on the plasma membrane expression of 232 clinical CF variants. Our results suggest that VX-121 provides a more robust rescue of variant expression for most CF variants (Fig. 1A). Nevertheless, we identify at least one CF-causing sequence variation (Y1032C) that directly perturbs the binding site in a manner that shifts its corrector selectivity (Fig. 3). This variant exhibits minimal difference in its proteostatic response to either corrector (Fig. 2). Moreover, functional rescue of Y1032C is selective toward combinations of modulators containing VX-445 rather than VX-121 (Fig. 4). Finally, to determine whether VX-121 is likely to have similar off-target effects relative to other clinically approved correctors, we used photo-crosslinking to determine whether it engages the known VX-445 off-target SCCPDH. Our results suggest structural modifications to VX-121 have abrogated its interaction with this enzyme (Fig. 5), which could potentially reduce the side effects of this compound relative to other treatments. Together, these investigations provide new insights into both the on- and off-target interactions of this CFTR modulator.

Though there are considerable structural differences between VX-121 and VX-445, our findings suggest CF variants that respond to one are likely to respond to the other. Nevertheless, our functional characterization of Y1032C and V1240G suggests there are likely to be few variants for which VX-445 offers superior proteostatic rescue. Interestingly, Y1032C has been annotated as a variant of varying clinical consequence (www.cftr2.org),(1) which may reflect its documented residual function (Fig. 4A).(26) Our new findings suggest that Y1032C is a VX-445 “super-responder,” as its functional rescue exceeds 90% of WT activity. Compared to VX-445 alone, combinations of VX-445 with VX-661 and/or VX-770 conferred no significant change to Y1032C activity. Moreover, VX-121 alone or in combination with VX-661 and VX-770 was ∼30% less effective than VX-445 (Fig. 4A). In contrast to Y1032C, V1240G has been classified as a CF-causing variant (www.cftr2.org) that retains minimal function (Fig. 4B).(26) Interestingly, our results show that VX-661 alone offers superior functional correction of V1240G relative to VX-445 or VX-121, regardless of whether VX-770 is included (Fig. 4B). Despite their distinct binding pockets, adding VX-445 does not enhance V1240G activity beyond what can be achieved with VX-661 alone (Fig. 4B). Despite the small amplitude of correction for V1240G conferred by VX-661, total channel activation is greater than 10% of WT; a benchmark that is widely considered to be the minimum CFTR rescue required to confer clinical improvement.(4, 9, 27, 28) Taken together, our findings indicate patients encoding Y1032C or V1240G could potentially experience greater benefit from Trikafta^TM^ or individual modulators rather than Alyftrek^TM^. Nevertheless, this possibility would require additional validation in primary human airway epithelia derived from CF patients encoding these specific variants.

The anomalous trends in the functional rescue of Y1032C and V1240G CFTR also serve as an important reminder that VX-445 acts as both a corrector and potentiator.(29–31) Given the allosteric aspects of CFTR modulator activity, it may be that VX-121 and VX-445 elicit non-equivalent potentiator activity in the context of certain variants. As Y1032C CFTR exhibits higher ion transport in the presence of VX-445 than VX-121 (Fig. 4A), it is possible that the direct structural interaction between the native Y1032 side chain and VX-121 participates in the allosteric propagation of conformational changes that give rise to potentiation. Unfortunately, we currently lack a robust means to quantitatively compare the extent to which these compounds correct the function of CFTR variants at scale using DMS. Higher-throughput transepithelial conductance assays may offer a more suitable approach for these purposes.(9) Additional investigations are needed to determine whether structural differences between VX-121 and VX-445 give rise to distinct effects on variant correction and potentiation.

Beyond differences in the direct effects of these compounds on CF channel variants, it is important to note that the indirect effects of these compounds that arise from off-target interactions also impact the utility of CFTR modulator therapies.(3) Using a modified VX-445 probe bearing a photo-activatable azide group, we recently showed that VX-445 forms a prominent off-target interaction with the SCCPDH enzyme associated with dopamine regulation,(25) which has also been reported as a promiscuous off-target for two other unrelated small molecules (WOBE437 and AX-1).(32) Given that the side effects of Trikafta^TM^ are broadly similar to those of Alyftrek^TM^, it seems unlikely that they are directly associated with variations in this specific off-target interaction. Nevertheless, divergent SCCPDH interactions could potentially lead to consequential differences in drug-drug interactions associated with these therapies. Additional investigations to comprehensively explore the off-targets of these CFTR modulators are needed to gain holistic insights into the manner in which these interactions are associated with unwanted clinical side effects.

## Materials and Methods

### Reagents

CFTR modulators were purchased from commercial providers. Compounds included elexacaftor (VX-445; #S885 Selleck Chemicals, Houston, TX, USA), tezacaftor (VX-661; #HY-15448 MedChemExpress, Monmouth Junction, NJ, USA), ivacaftor (VX-770; #HY-13017 MedChemExpress, Monmouth Junction, NJ, USA), vanzacaftor (VX-121; #E1701 Selleck Chemicals, Houston, TX, USA), deutivacaftor (VX-561; #HY-13017S MedChemExpress, Monmouth Junction, NJ, USA), amiloride (#A7410 Sigma-Aldrich, St. Louis, MO, USA), forskolin (#S2449 Selleck Chemicals, Houston, TX, USA), and CFTR Inhibitor172 (Inh172; #S7139 Selleck Chemicals, Houston, TX, USA). VU439 was prepared as previously described.(25)

### Cell Culture and Media

HEK293T cells were grown in high-glucose Dulbecco’s Modified Eagle’s Medium (DMEM) containing 10% fetal bovine serum (FBS), 1% penicillin/streptomycin mix, and 1% glutamine (Gibco, Grand Island, NY, USA). Parental Fischer rat thyroid (FRT) cells were a gift from Michael Welsh (University of Iowa, Iowa City, IA, USA). FRT cells were cultured in F12 Ham Coon’s modified nutrient mixture (#F6636 Sigma-Aldrich, St. Louis, MO, USA) supplemented with 2.68g sodium bicarbonate, 850μL 2N hydrochloric acid, and 5% fetal bovine serum (pH 7.3). Incubator conditions were set to 5% CO2, 95% O2, and 37°C as previously described. (10, 11)

### CFTR Expression Constructs

DMS studies were carried out using a previously described plasmid library containing 232 CF variants that are encoded in a CFTR cDNA containing a triple hemagglutinin (HA) epitope tag within the fourth extracellular loop (MSD-2 domain). This library also features an upstream Bxb1 recombination site to facilitate stable recombination and a downstream internal ribosome entry site (IRES)-GFP cassette, which serves as an expression reporter. This variant library was prepared via electroporation of electrocompetent NEB10β cells (New England Biolabs, Ipswich, MA, USA) and grown in LB broth overnight prior to purification using the ZymoPure endotoxin-free midiprep kit (Zymo Research, Irvine, CA, USA). Western blot and Ussing chamber measurements were carried out using an untagged CFTR expression construct in a pcDNA5 backbone that was generously provided by Eric Sorscher and Jeong Hong (Emory University, Atlanta, GA, USA). The V1240G and Y1032C variants were generated in this backbone by site-directed mutagenesis using the Q5 DNA polymerase and the KLD enzyme kit (New England Biolabs, Ipswich, MA, USA). The Bxb1 recombinase expression vector (pCAG-NLS-HA Bxb-1) that was used to generate recombinant cellular libraries was generously provided by Douglas Fowler (University of Washington, Seattle, WA, USA). The sequences of all plasmids used for these experiments were validated by whole plasmid sequencing (Plasmidsaurus Inc., Watterson Park, KY, USA).

### Western Blotting

Western blotting was carried out to detect transiently expressed CFTR proteins in HEK293T cells. Briefly, cells were seeded at 2×10^5^ cells/mL in 10 cm dishes and incubated for 24 hours prior to transfection using the calcium phosphate method. The transfection medium was then replaced with fresh medium after 16-18 hours. Cells were incubated for 1 hour before harvesting with trypsin-EDTA and 5 mL of medium. The cells were then evenly divided across a 6-well dish to ensure uniform transfection efficiency across the treatment conditions. Cells were then treated with either DMSO vehicle, 3µM VX-445, or 3µM VX-121 for 24 hours. Following treatment, cells were washed once with phosphate-buffered saline (PBS), then harvested using a cell scraper and pelleted at 1000 x g for 10 min in a microcentrifuge tube. Cells were then lysed by vortexing in 40 uL of 50 mM Tris (pH 7.5) containing 150 mM NaCl, 0.5% IGEPAL CA-630, and an EDTA-free Protease Inhibitor Cocktail (Roche Diagnostics, Indianapolis, IN, USA). Lysates were centrifuged at 12,000 × g for 10 minutes to remove any remaining insoluble material. The total protein concentration in each sample was then measured using a Pierce BCA Protein Assay kit (Thermo Scientific, Waltham, MA, USA) before normalizing the concentrations by dilution.

A 30µL SDS-PAGE sample was prepared for each lysate by adding concentrated Laemmli buffer containing 10 mM dithiothreitol (DTT) prior to warming the sample at 37°C for 15 minutes. Proteins were then separated using an 8% polyacrylamide SDS-PAGE gel. Proteins were transferred to a PVDF membrane for immunoblotting (Millipore Sigma, Burlington, MA, USA). The membrane was then blocked for 30 minutes with Tris-buffered saline (pH 7.5) with 0.1% Tween-20 (TBS-T) containing 5% non-fat dried milk (w/v). Blots were then incubated overnight at 4° C with a primary antibody wash solution consisting of TBS-T buffer containing 0.1% NaN_3_, 5% bovine serum albumin, and a 1: 1,000 dilution of an anti-CFTR 217 mouse primary antibody, which was provided by The Cystic Fibrosis Molecular/ Functional Measurement Core at the University of North Carolina, Chapel Hill (http://cftrantibodies.web.unc.edu/). Immunoblotted PVDF membranes were then washed three times in TBST and incubated with a secondary antibody wash solution consisting of a 1: 5,000 dilution of StarBright Blue 700-conjugated goat anti-mouse IgG secondary antibody (Biorad Labs, Hercules, CA, USA) in TBST containing 5% dried non-fat milk for 30 minutes at room temperature. The blot was then rinsed three additional times in TBST. A rhodamine-conjugated primary rabbit anti-tubulin antibody (Biorad Labs, Hercules, CA, USA) was included within the secondary antibody wash solutions to provide a secondary signal for the loading control. Blots were then imaged using a ChemiDoc MP Imaging System (Bio-Rad Labs, Hercules, CA, USA).

### Deep Mutational Scanning

Deep mutational scanning (DMS) experiments were carried out as previously described. Briefly, a genetically-modified HEK293T cell line bearing a genomic attP recombination site was used to prepare a pool of recombinant stable cells in which each cell inducibly expresses one of 232 known CF variants. Cells were grown at 37°C in an incubator containing 5% CO_2_ in Dulbecco’s Modified Eagle’s Medium (DMEM) containing 10% fetal bovine serum (FBS) and 1% penicillin/streptomycin mix (Gibco, Grand Island, NY, USA). Recombination was initiated by co-transfecting cells with a Bxb1 recombinase expression vector and a pooled CF variant library using Fugene 6 (Promega Corp., Madison, WI, USA). Cells were incubated at 33°C in an incubator containing 5% CO_2_ in Dulbecco’s Modified Eagle’s Medium (DMEM) with 10% fetal bovine serum, and 1% penicillin/streptomycin for 3 days post-transfection. Expression of CF variants was then initiated 24 hours post-transfection by adding 2 µg/ mL doxycycline to the growth medium. A Bigfoot Spectral Cell Sorter (Thermo Scientific, Waltham, MA, USA) was used to isolate recombinant cells expressing individual CF variants based on their characteristic gain in IRES-GFP expression via fluorescence-activated cell sorting. Recombinant cellular libraries were then expanded into 15-cm cell culture dishes and treated with the indicated concentrations of small molecules for 16 hours prior to harvesting with 0.25% Trypsin-EDTA (Gibco, Grand Island, NY, USA). Surface CFTR proteins on treated cellular libraries expressing CF variants were then marked using a DyLight 550-conjuaged with anti-HA antibody prior to fractionation of each treated library into four even quartiles (≥ 1 million cells/ fraction) according to relative surface immunostaining of expressed CF variants using a Bigfoot Spectral Cell Sorter (Thermo Scientific, Waltham, MA, USA). Each cellular isolate was then expanded in a 10 cm dish prior to harvesting and flash-freezing.

To quantify the relative abundance of each CF variant within each cellular isolate, we first extracted the genomic DNA (gDNA) from the frozen cell pellets using the DNEasy extraction kit (Qiagen Sciences Inc., Germantown, MD, USA). A previously described nested PCR approach was then used to create DNA amplicons bearing the region of the recombinant gDNA that contains unique molecular identifiers for the expressed CF variants. Amplicons generated for each cellular sub-population were then deeply sequenced (>7 million reads per amplicon) using a Cloudbreak Freestyle cell on an Aviti24 DNA sequencing system (Element Biosciences, San Diego, CA, USA). A previously described software package was then used to remove poor-quality reads, then estimate the relative surface immunostaining of each variant based on the relative abundance of each sequencing-based variant identification within each cellular fraction and the average immunostaining intensity of each fraction (https://github.com/schebachlab/RP-dms-analysis). Calculated average raw immunostaining intensities and scaled intensity values across three biological replicates can be found in Table S1.

To compare DMS-based responses to VX-121 to their corresponding VX-445 response, we compared these measurements to a recently described DMS data set collected in the presence of VX-445. Notably, previously described VX-445 treatment data were collected using a similar methodology but distinct instrumentation. Variations in the dynamic range of cell sorters used for these investigations complicate efforts to make direct, quantitative comparisons. Nevertheless, each drug-treated data set contains its own vehicle-treated intensity score for each variant. Therefore, to facilitate the identification of differences in drug response, we utilized the DMS-derived vehicle-treated variant intensity values, which should be identical, to derive an empirical coefficient that we used to scale the raw immunostaining intensities for each variant in the presence of one drug relative to its corresponding intensity in the presence of the other using the following equation:

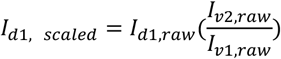

where *I*_d1, raw_ is the DMS-derived intensity for a variant in the presence of drug-one, *I*_v1, raw_ is the corresponding DMS-derived intensity for the vehicle-treated variant measurement associated with the drug-one data set, *I*_v2, raw_ is the DMS-derived intensity for the vehicle-treated variant measurement associated with the drug-two data set, and *I*_d1, scaled_ is the intensity for a variant in the presence of drug-one scaled relative to the corresponding intensities in the presence of drug two. To compare trends in expression across VX-121- and VX-445-treated CF variants, we used this formula to scale the intensities of VX-121-treated intensity values down to match the dynamic range of previously reported VX-445 measurements (Fig. 1E). Trends are similar when VX-445 measurements are instead scaled up to match the intensity range of VX-121 measurements (Figure S2).

### Flow Cytometry Measurements

The CFTR surface immunostaining intensities of individual CF variants were quantitatively compared by flow cytometry as previously described. Briefly, stable recombinant cells that inducibly express Y1032C or V1240G CFTRs were prepared using an HEK293T cell line bearing a genomic attP recombination site in the same manner used to prepare variant libraries described above, except that plasmid preps encoding single CF variants were co-transfected with the Bxb1 expression vector in place of the variant library. Cells were then treated and prepared for analysis in a similar manner as described above for DMS. Cellular fluorescence measurements were then recorded using a BD Fortessa Cell Analyzer (BD Biosciences, Franklin Lakes, NJ, USA) or a BigFoot Cell Sorter (Thermo Scientific, Waltham, MA, USA). Cellular fluorescence profiles were then analyzed using FlowJo software (Treestar Inc., Ashland, OR, USA).

### Computational Modeling

To model the binding of VX-121 within the CFTR transmembrane pocket, we first aligned VX-121 to the published VX-445 scaffold coordinates (PDB ID: 8EIG)(15) using the BioChemical Library (BCL) tool AlignToScaffold.(33) This was followed by the re-addition and randomization of hydrogen atoms to optimize local geometries. Following alignment, a structural ensemble of VX-121 conformers was generated using the BCL ConformerGenerator application.(34) The aligned VX-121 structure was neutralized, and a maximum of 2,000 iterations were performed to generate an ensemble of up to 250 low-energy conformers. Conformer diversity was ensured by using an RMSD comparer to filter redundant geometries and give a comprehensive representation of the ligand’s conformational space. To facilitate docking within the Rosetta software suite, ligand-specific topology and parameter files (.params) were generated using the molfile_to_params.py script. This translates the chemical properties and the pre-generated VX-121 conformer ensemble into a Rosetta-readable format and ensures accurate treatment of bond lengths, angles, and partial charges during the docking calculations.

For docking, we used an ensemble of active apo CFTR models (PDB ID: 6MSM) refined against the experimental cryo-EM density map using Rosetta as previously reported.(35, 36) The five lowest-scoring models by Rosetta Energy Units (REU) from this ensemble were used to dock VX-121 using RosettaLigand.(37) Our docking protocol consisted of an initial low-resolution perturbation followed by high-resolution refinement. Ligand-docked poses were first sampled using the Transform mover with a 5.0 Å box size and a 3.0 Å initial perturbation. High-resolution docking was performed using the HighResDocker (24 cycles) with a soft-repulsive scoring function followed by a FinalMinimizer stage using a standard ligand scoring function. To sample the binding interface, all side chains within 6.0 Å and backbone residues within 7.0 Å of the protein were allowed to move, with Cα restraints (0.3) applied to the backbone. Both the ligand_soft_rep and hard_rep score functions were customized with reweighted terms for electrostatic interactions (fa_elec = 0.42) and hydrogen bonding (hbond_bb_sc and hbond_sc = 1.3). We performed a total of 5,000 independent docking simulations. The resulting VX-121 docked models were ranked by the interface delta X score (i.e., interface energy), and the top 20 models were manually inspected. To investigate the reduced sensitivity of the Y1032C variant to VX-121, we used the MutateResidue mover to introduce the mutant using the same five active-apo state models described above. These were refined using the FastRelax mover to create an ensemble of 100 mutant structural models (20 per starting model). From this pool, the five lowest-energy mutant structures were selected prior to repeating the VX-121 docking protocol. All BCL/Rosetta XML scripts, command lines, and associated docking protocols reported in this study are available on the Meiler Lab GitHub repository (https://github.com/meilerlab/cftr-VX121).

### Short Circuit Current Measurements

FRT cells were grown and transfected on permeable filters to facilitate the formation of a monolayer with polarity. Cells were transfected with wild-type (WT) or mutant CFTR cDNA (0.5 μg) based on an established protocol. (10, 11, 37, 38) In short, cells were seeded on 0.33cm^2^ permeable transwells at 1.5 x 10^5^ cells/well. The following day, plasmids were transfected onto the apical surface of FRT monolayers using the Lipofectamine 3000 kit (Invitrogen, Carlsbad, CA, USA) and 1X Opti-MEM Reduced Serum (Gibco, Grand Island, NY, USA). Cells were then incubated in the transfection medium for 24 hours prior to aspiration and the formation of a cellular air-liquid interface. Cells were treated basolaterally under chronic incubation with CFTR modulators (5 μM per drug) for 48 hours prior to functional assessment. Cells were mounted in P2300 Ussing chambers (MC8 Apparatus, Physiologic Instruments, Venice, FL, USA) and analyzed for short-circuit current electrophysiology as previously described.(10, 11, 38, 39) Briefly, cells were equilibrated for 5-10 minutes with basolateral regular Ringer and apical low-chloride Ringer, after which the following antagonists or agonists were acutely delivered to the apical and/or basolateral surfaces: amiloride (100 μM; sodium channel inhibitor), forskolin (10 μM; CFTR activator via stimulation of cAMP-dependent Protein Kinase A phosphorylation), VX-770 (5 μM; CFTR gating potentiator), and Inh172 (10 μM; CFTR-specific inhibitor). CFTR-mediated transepithelial ion transport was measured as the difference (Δ) from baseline current to the highest or lowest value of a stable plateau achieved for 5-10 minutes.

### Photo-Crosslinking and Immunoprecipitation

To compare the off-target interactions of VX-121 and VX-445, HEK293T cells were first seeded into 10 cm cell culture dishes at a density of 1.5 × 10^6^ cells/ plate 24 hours prior to transfection. Cells were then transfected with 5 µg of SCCPDH-Myc expression construct or a control GFP expression construct using the calcium phosphate method. Cells were incubated for 16-18 hours, then washed twice with PBS prior to replacing the medium. Cells were then incubated for one hour before being split into 6-well dishes coated with poly-D-lysine hydrobromide according to the manufacturer’s protocol (Sigma Aldrich, St. Louis, MO, USA). SCCPDH-Myc expressing cells were pre-treated with any unlabeled competing molecules for two hours before treatment with 3 µM VU-439 or the DMSO vehicle control for 18 hours. Cells treated with VU439 treatment were switched into Opti-MEM medium (Thermo Fisher, Waltham, MA, USA) prior to the initiation of photo crosslinking with 368 nm light for 2 minutes (600 mJ total dosage). Cells were then washed with cold PBS and lysed by adding 200 µL Radioimmunoprecipitation assay (RIPA) buffer consisting of 50 mM Tris (pH 7.5) containing 150 mM NaCl, 1% Triton X-100 (v/v), 0.5% sodium deoxycholate (w/v), and 0.1% SDS (w/v) supplemented with complete EDTA-free protease inhibitor (Sigma Aldrich, St. Louis, MO, USA) to each well then rocking the plate at 4°C for 15 minutes. The lysate was cleared by centrifuging at 12,000 × g for 10 minutes to remove any remaining insoluble material prior to the quantification of total protein concentration in each sample using a Pierce BCA Protein Assay kit (Thermo Scientific, Waltham, MA, USA). To biotinylate cross-linked proteins, lysates were diluted to a total protein concentration of 1 mg/mL in 170 µL prior to the addition of 30 µL Click master mix to achieve a final concentration of 100 μM TAMRA-Azide-PEG-Desthiobiotin (BroadPharm, San Diego, CA, USA), 1.6 mM BTTAA (Vector Labs, Newark, CA, USA), 5 mM sodium ascorbate, and 0.8 mM Cu_2_SO_4_. Click-modified proteins within lysates were precipitated with 3:1:3 methanol: chloroform: water, then washed twice with methanol. After drying at room temperature, protein pellets were resuspended in 100 µL of 6M urea containing 1% SDS (w/v), then diluted in PBS to a final volume of 1100 µL. To immunoprecipitate biotinylated proteins, 100 µL of prewashed, high-capacity streptavidin resin (Thermo Fisher, Waltham, MA, USA) was added to each sample, then rotated at room temperature for 24 hours. Beads were washed with four consecutive PBS solutions containing 1% SDS (w/v), 4M urea, 1 M NaCl, and 1% SDS (w/v) before the remaining liquid was removed using a 30 G needle. Beads were then frozen at -80 °C overnight prior to eluting twice with 100 µL of an elution buffer containing 50 mM biotin and 1% SDS (w/v) at 95 °C for 5 minutes.

## Acknowledgements

We thank Phillip San Miguel at the Bindley Bioscience Center Genomics Facility for experimental support. This research was supported by grants from the National Institutes of Health (NIH) (R01HL167046 to JPS, LP, KEO, and JM; T32GM149371 to JAO and CDC). MP was supported by the Vanderbilt SyBBURE Searle Program.

## Data Availability

Code for the analysis of sequencing data for deep mutational scanning experiments can be found on the Schlebach Lab GitHub page (https://github.com/schebachlab/RP-dms-analysis). Illumina sequencing data is freely available as an NCBI BioProject (SAMN60458616). DMS measurements are included as Excel files in the Supplementary Materials. Structural modeling data and any associated code can also be accessed via GitHub (https://github.com/meilerlab/cftr-VX121). All other experimental data have been deposited in a Mendeley data directory (doi: 10.17632/3gxywvsxns.1).

