## Supplemental Materials for "Profiling the CFTR Variant Selectivity and Off-Target Interactions of VX-121"

‡ Authors contributed equally

*Corresponding Authors: jschleba (at) purdue.edu, lars.plate (at) vanderbilt.edu, kolive3 (at) emory.edu,

Contents:

-Figure S1
-Figure S2
-Table S1

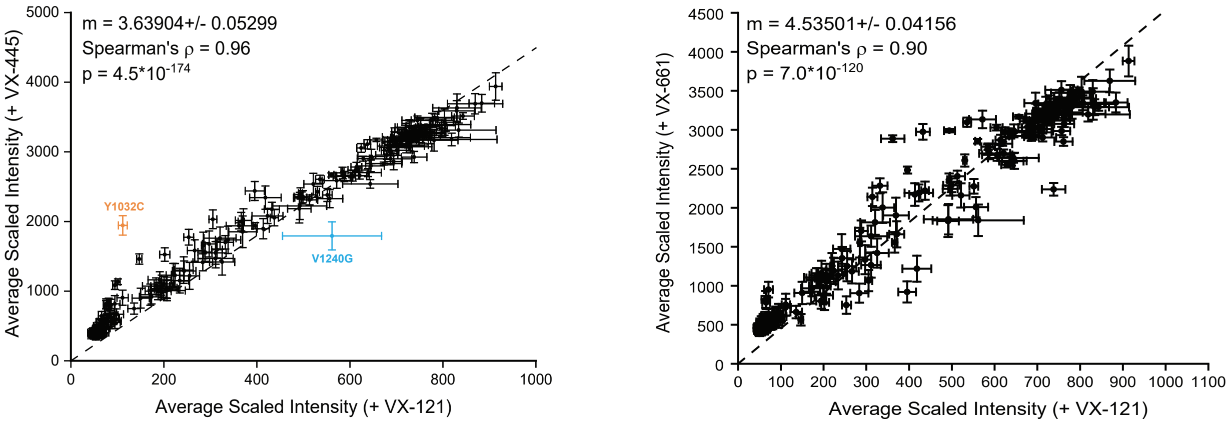


**Figure S1. Comparison of Surface Immunostaining Intensities of CF Variants in the Presence of VX-121, VX-445, and VX-661.** The surface immunostaining intensities of 232 CF variants were determined in the presence of 3 μM VX-121 by deep mutational scanning. Intensity values in the presence of VX-121 are plotted against the corresponding intensities in the presence of A) 3 μM VX-445 or B) 3 μM VX-661 Measurements represent the average of 3 biological replicates. The error bars represent Standard Error. A linear fit line (black dashes) is shown for reference and the fitted values of the slope (*m*) is shown along with the and Spearman's Correlation coefficient (ρ), for reference.

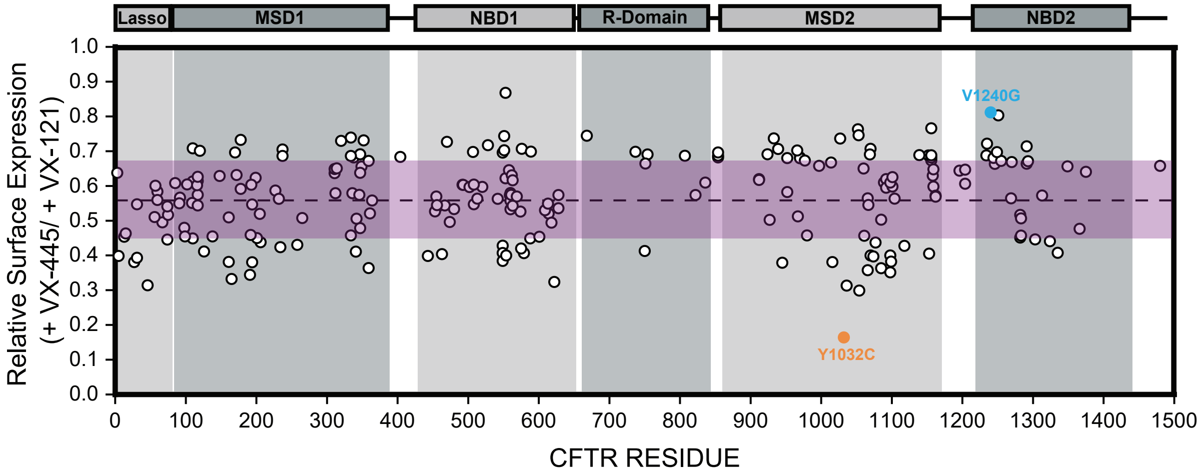


**Figure S2. Alternative Scaling for CF Variants VX-121/ VX-445 Response Ratio.** CFTR surface immunostaining intensities for each variant in the presence vehicle were used to scale the corresponding intensity values in the presence of 3 μM VX-121 or 3 μM VX-445. The ratio of the VX-445-treated intensity to the corresponding VX-121-treated intensity for each variant is plotted against the position of the mutated residues within the CFTR sequence. The boundaries of the lasso motif, membrane spanning domain 1 (MSD1), nucleotide binding domain 1 (NBD1), the R-domain, membrane spanning domain 2 (MSD2), and nucleotide binding domain 2 (NBD2) are highlighted in gray for reference, and the purple stripe reflects values within one standard deviation of the mean. Trends are similar to the reciprocol VX-121/ VX-445 ratios that are plotted in Figure 1E.

### Table S1. Deep Mutational Scanning Measurements of CF Variant Immunostaining Intensities.

| CFTR Sub-Domain | Variant | Amino Acid Substitution | Vehicle Control -Average Scaled Intensity (For VX-121) | VX-121 Average Scaled Intensity | Up-Scaled VX-121 Average Scaled Intensity | Vehicle Control - Average Scaled Intensity (For VX-445) | VX-445 Average Scaled Intensity | Vehicle Control - Average Scaled Intensity (For VX-121+VX-661) | VX-121+VX-661 Average Scaled Intensity |
|---|---|---|---|---|---|---|---|---|---|
| Lasso | R3W | 7A>T | 315.2 | 646.2 | 1871.0 | 912.6 | 2934.1 | 741.0 | 3352.7 |
| Lasso | P5L | 14C>T | 105.6 | 395.4 | 971.5 | 259.6 | 2438.3 | 123.7 | 1959.3 |
| Lasso | S13F | 38C>T | 56.1 | 77.5 | 279.6 | 202.4 | 616.2 | 63.1 | 579.5 |
| Lasso | L15P | 44T>C | 60.9 | 71.1 | 248.0 | 212.3 | 535.3 | 79.8 | 507.8 |
| Lasso | G27R | 79G>A | 55.4 | 66.2 | 245.2 | 205.3 | 644.5 | 64.9 | 517.5 |
| Lasso | R31C | 91C>T | 215.2 | 492.2 | 1216.1 | 531.7 | 2223.6 | 448.6 | 2135.9 |
| Lasso | R31L | 92G>T | 159.8 | 418.3 | 920.7 | 351.8 | 2342.9 | 179.1 | 2037.7 |
| Lasso | A46D | 137C>A | 70.0 | 147.1 | 458.7 | 218.2 | 1460.9 | 67.7 | 1017.9 |
| Lasso | E56K | 166G>A | 59.3 | 64.9 | 227.3 | 207.7 | 445.6 | 57.3 | 821.3 |
| Lasso | W57G | 169T>G | 44.7 | 52.8 | 235.6 | 199.6 | 392.3 | 71.3 | 303.6 |
| Lasso | E60X | 178G>T | 48.5 | 53.7 | 229.1 | 206.8 | 392.5 | 71.2 | 363.7 |
| Lasso | E60K | 178G>A | 47.0 | 51.4 | 217.0 | 198.1 | 387.9 | 63.7 | 466.7 |
| Lasso | P67L | 200C>T | 63.1 | 65.8 | 221.5 | 212.4 | 447.5 | 77.3 | 733.8 |
| Lasso | R74W | 220C>T | 214.5 | 288.3 | 695.9 | 517.8 | 1562.0 | 385.8 | 1513.6 |
| Lasso | R74Q | 221G>A | 288.2 | 432.2 | 1175.8 | 784.1 | 2174.8 | 626.3 | 3242.0 |
| Lasso | R75Q | 224G>A | 274.2 | 361.6 | 1001.1 | 759.2 | 1937.8 | 589.3 | 2898.0 |
| TMD1 | G85E | 254G>A | 45.3 | 52.7 | 233.3 | 200.2 | 383.8 | 62.9 | 306.7 |
| TMD1 | G91R | 271G>A | 51.1 | 59.1 | 224.9 | 194.5 | 409.0 | 71.3 | 654.5 |
| TMD1 | E92K | 274G>A | 50.3 | 56.6 | 228.9 | 203.7 | 404.6 | 65.4 | 338.2 |
| TMD1 | Q98R | 293A>G | 63.7 | 71.1 | 234.9 | 210.3 | 490.2 | 71.9 | 915.1 |
| TMD1 | P99L | 296C>T | 188.9 | 283.8 | 642.7 | 427.8 | 1414.8 | 286.7 | 1083.5 |
| TMD1 | L102R | 305T>G | 43.2 | 50.0 | 230.2 | 199.1 | 381.1 | 57.4 | 416.5 |
| TMD1 | Y109N | 325T>A | 275.6 | 396.3 | 1068.0 | 742.7 | 1938.7 | 695.2 | 2980.0 |
| TMD1 | D110Y | 328G>T | 217.5 | 314.4 | 715.9 | 495.4 | 1594.4 | 370.4 | 2264.9 |
| TMD1 | D110E | 330C>A | 441.2 | 756.4 | 2291.4 | 1336.6 | 3237.4 | 1188.1 | 3081.6 |
| TMD1 | D110H | 328G>C | 357.3 | 535.7 | 1596.4 | 1064.9 | 2606.2 | 868.5 | 2757.1 |
| TMD1 | E116K | 346G>A | 344.5 | 494.3 | 1409.2 | 982.0 | 2342.6 | 829.9 | 2991.9 |
| TMD1 | R117C | 349C>T | 322.4 | 513.1 | 1326.1 | 833.3 | 2310.2 | 679.5 | 2418.5 |
| TMD1 | R117H | 350G>A | 362.4 | 560.1 | 1638.1 | 1059.8 | 2668.3 | 825.4 | 2722.5 |
| TMD1 | R117P | 350G>C | 356.9 | 606.2 | 1662.6 | 978.8 | 2655.8 | 905.9 | 3200.9 |
| TMD1 | R117L | 350G>T | 285.2 | 422.7 | 1130.1 | 762.4 | 2079.8 | 621.4 | 2570.3 |
| TMD1 | A120T | 358G>A | 431.6 | 747.8 | 2255.2 | 1301.5 | 3217.0 | 1116.2 | 3501.5 |
| TMD1 | G126D | 377G>A | 158.2 | 242.5 | 535.9 | 349.5 | 1302.7 | 193.7 | 1791.6 |
| TMD1 | L138ins | 413_415dupTAC | 53.5 | 58.8 | 221.1 | 201.2 | 486.5 | 65.6 | 565.4 |
| TMD1 | I148T | 443T>C | 368.9 | 571.6 | 1625.7 | 1049.1 | 2588.0 | 886.0 | 2830.0 |
| TMD1 | Y161D | 481T>G | 48.8 | 53.3 | 217.6 | 199.3 | 427.2 | 59.2 | 476.2 |
| TMD1 | Y161C | 482A>C | 60.0 | 83.0 | 285.4 | 206.4 | 748.6 | 61.7 | 907.8 |
| TMD1 | L165S | 494T>C | 56.4 | 75.8 | 267.7 | 199.1 | 806.1 | 71.6 | 505.5 |
| TMD1 | R170H | 509G>A | 401.7 | 679.3 | 2055.6 | 1215.7 | 2953.4 | 1077.9 | 3350.9 |
| TMD1 | L172I | 514C>A | 362.3 | 637.5 | 1963.0 | 1115.7 | 3109.0 | 868.3 | 3209.2 |
| TMD1 | G178R | 532G>A | 375.8 | 585.0 | 1570.3 | 1008.9 | 2655.4 | 781.9 | 2691.3 |
| TMD1 | G178E | 533G>A | 444.6 | 806.2 | 2483.0 | 1369.1 | 3391.7 | 1214.5 | 3472.2 |

| | | | | | | | | | |
|---|---|---|---|---|---|---|---|---|---|
| TMD1 | F191V | 571T>G | 89.6 | 201.9 | 524.8 | 233.0 | 1525.5 | 93.6 | 1665.8 |
| TMD1 | D192G | 575A>G | 78.0 | 253.4 | 814.4 | 250.6 | 1774.5 | 103.9 | 1299.2 |
| TMD1 | E193K | 577G>A | 303.8 | 530.4 | 1463.0 | 838.1 | 2425.6 | 685.2 | 3174.1 |
| TMD1 | G194R | 580G>A | 287.6 | 521.2 | 1386.5 | 765.0 | 2537.4 | 647.4 | 2809.7 |
| TMD1 | G194V | 581G>T | 142.0 | 220.0 | 469.0 | 302.6 | 1233.5 | 192.9 | 1826.9 |
| TMD1 | H199Y | 595C>T | 48.5 | 58.7 | 243.7 | 201.4 | 391.2 | 74.9 | 391.1 |
| TMD1 | V201M | 601G>A | 215.3 | 331.9 | 765.4 | 496.5 | 1701.1 | 380.2 | 2362.5 |
| TMD1 | P205S | 613C>T | 50.3 | 55.6 | 226.7 | 205.1 | 436.6 | 57.5 | 388.7 |
| TMD1 | L206W | 617T>G | 54.0 | 64.9 | 243.2 | 202.5 | 555.6 | 69.0 | 561.4 |
| TMD1 | L227R | 680T>G | 47.4 | 55.7 | 235.3 | 200.3 | 401.2 | 80.9 | 411.4 |
| TMD1 | V232D | 695T>A | 51.3 | 63.6 | 250.7 | 202.2 | 445.8 | 64.9 | 471.0 |
| TMD1 | A234D | 701C>A | 200.5 | 311.0 | 656.2 | 423.1 | 1549.3 | 325.0 | 1802.9 |
| TMD1 | Q237E | 709C>G | 428.4 | 762.7 | 2312.7 | 1299.0 | 3282.1 | 1118.6 | 3353.9 |
| TMD1 | Q237H | 711G>C | 441.2 | 788.1 | 2391.9 | 1339.2 | 3486.7 | 1155.7 | 3595.3 |
| TMD1 | R258G | 772A>G | 100.8 | 136.0 | 322.1 | 238.7 | 749.1 | 91.9 | 889.9 |
| TMD1 | M265R | 794T>G | 72.6 | 94.5 | 289.8 | 222.7 | 571.2 | 61.6 | 602.8 |
| TMD1 | F311L | 933C>G | 411.2 | 686.0 | 1999.7 | 1198.8 | 3140.8 | 940.6 | 2962.5 |
| TMD1 | F311L | 933C>A | 413.8 | 712.4 | 2054.9 | 1193.7 | 3168.7 | 961.4 | 3296.1 |
| TMD1 | F312DEL | 935_937del | 267.2 | 556.5 | 1347.5 | 647.1 | 2328.7 | 574.0 | 2787.1 |
| TMD1 | G314E | 941G>A | 325.3 | 639.1 | 1757.2 | 894.4 | 2702.6 | 761.0 | 3062.9 |
| TMD1 | L320V | 958T>G | 450.7 | 803.6 | 2498.1 | 1401.0 | 3425.2 | 1190.4 | 3540.4 |
| TMD1 | R334W | 1000C>T | 461.9 | 797.6 | 2467.8 | 1429.3 | 3338.9 | 1184.9 | 3227.9 |
| TMD1 | R334L | 1001G>T | 73.9 | 92.8 | 275.3 | 219.2 | 602.2 | 89.3 | 510.2 |
| TMD1 | R334Q | 1001G>A | 441.0 | 756.9 | 2248.4 | 1310.1 | 3276.8 | 1095.5 | 3304.7 |
| TMD1 | I336K | 1007T>A | 54.1 | 72.3 | 271.3 | 202.9 | 468.7 | 70.3 | 562.9 |
| TMD1 | T338I | 1013C>T | 407.9 | 721.3 | 2111.6 | 1194.0 | 3098.1 | 985.6 | 2767.5 |
| TMD1 | S341P | 1021T>C | 166.9 | 243.5 | 529.8 | 363.1 | 1289.8 | 230.2 | 1586.8 |
| TMD1 | R342W | 1054C>T | 214.1 | 311.2 | 742.9 | 511.1 | 1469.8 | 355.1 | 1974.5 |
| TMD1 | L346P | 1037T>C | 43.5 | 51.4 | 221.9 | 188.0 | 386.6 | 64.1 | 559.2 |
| TMD1 | R347H | 1040G>A | 484.7 | 883.6 | 2553.5 | 1400.7 | 3697.1 | 1218.9 | 3036.6 |
| TMD1 | R347P | 1040G>C | 90.5 | 200.5 | 528.4 | 238.5 | 1104.3 | 134.2 | 1249.2 |
| TMD1 | R347P | 1040G>T | 339.5 | 738.1 | 1865.3 | 858.1 | 2925.4 | 849.1 | 3110.9 |
| TMD1 | A349V | 1046C>T | 358.4 | 604.7 | 1746.9 | 1035.5 | 2663.9 | 844.6 | 2642.9 |
| TMD1 | R352Q | 1055G>A | 467.6 | 843.1 | 2568.3 | 1424.3 | 3514.9 | 1212.0 | 3118.8 |
| TMD1 | Q359K/T360K | 1075-1079C>A | 151.5 | 266.3 | 575.6 | 327.4 | 1582.8 | 200.5 | 1564.8 |
| TMD1 | Q359R | 1076A>G | 403.2 | 689.0 | 2023.5 | 1184.1 | 3011.4 | 1069.9 | 3464.4 |
| TMD1 | W361R | 1081T>C | 52.0 | 59.6 | 222.6 | 194.2 | 427.5 | 68.6 | 418.0 |
| TMD1 | S364P | 1090T>C | 51.8 | 58.6 | 236.1 | 208.8 | 423.3 | 53.9 | 397.7 |
| XXX | G404R | 1210G>C | 416.1 | 695.8 | 2194.1 | 1312.0 | 3212.0 | 1058.9 | 3405.7 |
| NBD1 | D443Y | 1327G>T | 165.5 | 195.3 | 426.6 | 361.5 | 1070.9 | 236.3 | 1450.2 |
| NBD1 | L453S | 1358T>C | 54.5 | 58.7 | 262.9 | 243.8 | 499.1 | 63.7 | 424.6 |
| NBD1 | A455E | 1364C>A | 51.4 | 59.4 | 227.8 | 197.3 | 400.0 | 71.4 | 343.4 |
| NBD1 | V456A | 1367T>C | 48.5 | 52.6 | 214.5 | 197.8 | 398.1 | 103.6 | 424.0 |
| NBD1 | V456F | 1366G>T | 48.8 | 53.4 | 222.3 | 203.4 | 408.0 | 73.0 | 386.3 |
| NBD1 | G463D | 1388G>T | 54.2 | 79.8 | 294.9 | 200.2 | 731.1 | 58.2 | 585.6 |
| NBD1 | L467P | 1400T>C | 47.2 | 49.9 | 199.6 | 188.9 | 367.3 | 63.0 | 416.2 |

| | | | | | | | | | |
|---|---|---|---|---|---|---|---|---|---|
| NBD1 | M470V | 1408A>G | 417.0 | 645.1 | 2033.6 | 1314.6 | 2798.3 | 1038.3 | 2354.1 |
| NBD1 | E474K | 1420G>A | 48.8 | 55.9 | 224.3 | 195.8 | 452.4 | 70.7 | 437.5 |
| NBD1 | G480S | 1438G>T | 53.5 | 60.5 | 233.2 | 206.2 | 437.5 | 69.2 | 420.0 |
| NBD1 | S492F | 1475C>T | 50.3 | 58.1 | 225.8 | 195.5 | 374.3 | 73.5 | 345.7 |
| NBD1 | Q493X | 1477C>T | 44.3 | 51.8 | 231.5 | 198.0 | 384.3 | 64.3 | 360.0 |
| NBD1 | I502T | 1505T>C | 50.1 | 56.3 | 224.0 | 199.5 | 375.4 | 67.4 | 329.6 |
| NBD1 | I507del | 1519_1521delATC | 48.3 | 61.9 | 254.7 | 198.8 | 365.2 | 64.0 | 379.4 |
| NBD1 | F508del | 1521_1523delCTT | 50.3 | 57.9 | 224.5 | 195.1 | 410.3 | 73.1 | 375.1 |
| NBD1 | F508C | 1523T>G | 343.5 | 624.3 | 1844.4 | 1014.7 | 3055.5 | 876.2 | 3414.7 |
| NBD1 | D513G | 1538A>G | 49.2 | 56.2 | 227.4 | 198.7 | 403.7 | 62.8 | 400.3 |
| NBD1 | V520F | 1558G>T | 46.6 | 52.4 | 226.5 | 201.4 | 379.6 | 68.3 | 364.7 |
| NBD1 | E528E | 1584G>A | 436.4 | 761.3 | 2352.7 | 1348.7 | 3280.6 | 1095.0 | 3400.6 |
| NBD1 | G542X | 1624G>T | 41.5 | 45.9 | 226.7 | 205.2 | 402.9 | 64.8 | 407.7 |
| NBD1 | S549R | 1645A>C | 188.7 | 205.4 | 439.3 | 403.7 | 1027.0 | 287.9 | 1177.7 |
| NBD1 | S549N | 1646G>A | 442.8 | 741.6 | 2244.1 | 1340.0 | 3226.6 | 1114.2 | 2879.0 |
| NBD1 | S549R | 1647T>A | 174.2 | 183.3 | 436.8 | 414.9 | 1072.9 | 267.6 | 1136.7 |
| NBD1 | S549R | 1647T>G | 187.6 | 194.2 | 398.1 | 384.7 | 1036.1 | 307.5 | 1027.9 |
| NBD1 | G551S | 1651G>A | 470.1 | 869.4 | 2741.6 | 1482.6 | 3690.0 | 1245.6 | 3317.3 |
| NBD1 | G551D | 1652G>A | 470.6 | 829.5 | 2519.1 | 1429.1 | 3587.9 | 1149.3 | 3232.5 |
| NBD1 | R553X | 1657C>T | 44.3 | 54.7 | 250.3 | 203.1 | 402.5 | 70.3 | 357.6 |
| NBD1 | R553N | 1658G>A | 395.9 | 818.7 | 2757.7 | 1333.6 | 3178.8 | 1173.8 | 3457.7 |
| NBD1 | A554E | (1661C>A) | 155.3 | 173.3 | 367.2 | 329.2 | 918.2 | 219.9 | 1249.9 |
| NBD1 | L558S | 1673T>C | 50.4 | 61.6 | 235.4 | 192.7 | 364.8 | 69.0 | 328.2 |
| NBD1 | A559T | 1675G>A | 52.5 | 60.0 | 228.1 | 199.5 | 397.9 | 83.6 | 326.8 |
| NBD1 | R560K | 1679G>A | 44.7 | 48.7 | 213.4 | 195.8 | 368.4 | 66.5 | 342.2 |
| NBD1 | R560T | 1679G>C | 50.9 | 55.4 | 220.2 | 202.2 | 395.6 | 88.6 | 371.9 |
| NBD1 | R560S | 1680A>C | 49.3 | 52.4 | 203.7 | 191.5 | 382.4 | 65.0 | 460.6 |
| NBD1 | A561E | 1682C>A | 47.0 | 52.3 | 215.9 | 193.9 | 376.0 | 73.9 | 349.4 |
| NBD1 | V562I | 1684G>A | 372.6 | 588.0 | 1668.2 | 1057.0 | 2646.5 | 857.1 | 2398.1 |
| NBD1 | Y563N | 1687T>A | 45.4 | 52.7 | 224.0 | 193.2 | 363.7 | 60.0 | 393.1 |
| NBD1 | Y563D | 1687T>G | 45.8 | 50.0 | 210.6 | 192.9 | 388.1 | 61.2 | 405.0 |
| NBD1 | Y569D | 1705T>G | 48.0 | 54.2 | 232.4 | 205.6 | 408.2 | 68.7 | 317.7 |
| NBD1 | P574H | 1721C>A | 53.0 | 60.5 | 231.5 | 202.9 | 440.3 | 66.5 | 359.9 |
| NBD1 | F575Y | 1724T>A | 82.8 | 86.6 | 238.0 | 227.5 | 566.8 | 65.9 | 502.7 |
| NBD1 | G576A | 1727G>C | 432.4 | 731.3 | 2367.8 | 1400.0 | 3352.2 | 1119.8 | 2827.2 |
| NBD1 | D579G | 1736A>G | 183.5 | 187.6 | 406.5 | 397.5 | 1000.0 | 265.0 | 1062.2 |
| NBD1 | E588V | 1763A>T | 190.4 | 206.6 | 452.9 | 417.2 | 1008.8 | 306.4 | 1234.4 |
| NBD1 | S589T | 1766G>C | 398.2 | 672.7 | 1977.4 | 1170.4 | 2831.8 | 1017.2 | 3066.6 |
| NBD1 | I601F | 1801A>T | 84.0 | 88.7 | 249.3 | 236.1 | 549.8 | 83.2 | 547.5 |
| NBD1 | H609R | 1826A>G | 48.1 | 53.3 | 223.2 | 201.4 | 421.3 | 61.9 | 321.8 |
| NBD1 | A613T | 1837G>A | 53.9 | 58.7 | 226.7 | 208.1 | 413.0 | 69.6 | 487.2 |
| NBD1 | D614G | 1841A>G | 58.6 | 64.8 | 233.3 | 211.1 | 448.3 | 77.0 | 396.7 |
| NBD1 | I618T | 1853T>C | 60.2 | 67.1 | 232.6 | 208.8 | 470.6 | 51.8 | 416.3 |
| NBD1 | G622D | 1865G>A | 88.2 | 111.2 | 292.8 | 232.4 | 905.6 | 89.7 | 1056.1 |
| NBD1 | G628R | 1882G>A | 54.6 | 60.4 | 224.4 | 202.9 | 418.9 | 65.9 | 361.3 |
| NBD1 | G628R | 1882G>C | 57.6 | 67.0 | 228.1 | 196.3 | 397.8 | 75.0 | 403.5 |

| | | | | | | | | | |
|---|---|---|---|---|---|---|---|---|---|
| R-Domain | R668C | 2002C>T | 385.7 | 687.5 | 2157.4 | 1210.5 | 2900.0 | 983.3 | 2654.3 |
| R-Domain | S737F | 2210C>T | 443.4 | 742.0 | 2283.8 | 1364.7 | 3272.8 | 1087.9 | 3470.8 |
| R-Domain | P750L | 2249C>T | 160.3 | 223.9 | 506.8 | 362.9 | 1228.0 | 240.3 | 1280.5 |
| R-Domain | R751L | 2252G>T | 411.5 | 686.1 | 1999.5 | 1199.2 | 3009.8 | 1051.4 | 3209.0 |
| R-Domain | V754M | 2260G>A | 433.7 | 725.5 | 2201.5 | 1316.1 | 3189.0 | 1073.5 | 2955.2 |
| R-Domain | I807M | 2374C>G | 436.8 | 733.4 | 2231.6 | 1329.2 | 3250.5 | 1115.9 | 3462.4 |
| R-Domain | E822K | 2464G>A | 249.3 | 491.1 | 1253.1 | 636.2 | 2184.0 | 533.8 | 2613.0 |
| R-Domain | D836Y | 2506G>T | 370.7 | 583.9 | 1654.2 | 1050.2 | 2714.7 | 880.0 | 2940.0 |
| TMD2 | T854T | 2562T>A | 433.3 | 733.5 | 2190.8 | 1294.4 | 3212.5 | 1088.0 | 2943.4 |
| TMD2 | T854T | 2562T>C | 425.3 | 718.9 | 2246.9 | 1329.4 | 3270.3 | 1063.5 | 3026.5 |
| TMD2 | T854T | 2562T>G | 423.3 | 724.3 | 2202.3 | 1287.0 | 3164.3 | 984.6 | 2855.7 |
| TMD2 | S912X | 2735C>A | 47.6 | 56.3 | 239.3 | 202.0 | 386.5 | 59.3 | 333.5 |
| TMD2 | S912L | 2735C>T | 322.3 | 499.0 | 1453.3 | 938.8 | 2355.8 | 772.4 | 2700.5 |
| TMD2 | D924N | 2770G>A | 434.4 | 830.3 | 2508.3 | 1312.3 | 3630.0 | 1167.4 | 3778.7 |
| TMD2 | L927P | 2780T>C | 244.7 | 320.6 | 884.2 | 674.9 | 1760.4 | 448.9 | 1961.3 |
| TMD2 | R933G | 2797A>G | 508.9 | 913.5 | 2901.7 | 1616.6 | 3938.5 | 1378.4 | 3901.7 |
| TMD2 | H939R | 2816A>G | 407.4 | 782.1 | 2466.8 | 1285.0 | 3494.1 | 1092.8 | 3314.0 |
| TMD2 | S945L | 2834C>T | 123.2 | 191.2 | 440.3 | 283.7 | 1162.9 | 145.6 | 1073.7 |
| TMD2 | M952T | 2855T>C | 415.2 | 708.6 | 2118.3 | 1241.2 | 3115.0 | 1066.4 | 3444.4 |
| TMD2 | M952I | 2856G>A | 340.7 | 628.8 | 1690.3 | 915.9 | 2902.7 | 766.8 | 3072.3 |
| TMD2 | Q966= | 2988G>A | 425.2 | 720.3 | 2243.3 | 1324.2 | 3196.2 | 1090.7 | 2969.5 |
| TMD2 | L967S | 2900T>C | 283.5 | 369.3 | 948.6 | 728.2 | 1851.9 | 564.9 | 1950.3 |
| TMD2 | G970R | 2908G>C | 449.5 | 727.3 | 2197.9 | 1358.5 | 3224.6 | 1163.3 | 2968.1 |
| TMD2 | G970D | 2909G>A | 385.6 | 618.5 | 1860.7 | 1159.9 | 2738.5 | 983.2 | 2637.1 |
| TMD2 | S977F | 2930C>T | 393.5 | 711.8 | 2203.6 | 1218.2 | 3276.5 | 922.4 | 2992.2 |
| TMD2 | I980K | 2939T>A | 72.3 | 98.3 | 305.1 | 224.4 | 667.3 | 64.3 | 653.7 |
| TMD2 | L997F | 2991G>C | 426.1 | 745.3 | 2268.4 | 1297.0 | 3451.1 | 1034.7 | 3177.2 |
| TMD2 | Y1014C | 3041A>G | 383.5 | 692.4 | 2056.6 | 1139.1 | 3082.9 | 980.2 | 3301.6 |
| TMD2 | F1016S | 3047T>C | 168.1 | 297.3 | 638.2 | 360.9 | 1677.5 | 279.8 | 2221.3 |
| TMD2 | I1027T | 3080T>C | 354.9 | 643.9 | 1869.2 | 1030.2 | 2539.9 | 821.7 | 2347.2 |
| TMD2 | Y1032C | 3095A>G | 82.0 | 111.6 | 319.0 | 234.5 | 1944.2 | 107.4 | 795.7 |
| TMD2 | T1036N | 3107C>A | 60.9 | 101.2 | 355.1 | 213.9 | 1135.0 | 72.8 | 600.0 |
| TMD2 | F1052V | 3154T>G | 395.0 | 773.5 | 2444.4 | 1248.4 | 3204.5 | 1056.4 | 2959.3 |
| TMD2 | T1053I | 3158C>T | 414.0 | 728.8 | 2292.3 | 1302.3 | 3074.0 | 1050.1 | 3094.2 |
| TMD2 | H1054D | 3160C>G | 60.4 | 94.7 | 324.2 | 206.7 | 1086.2 | 69.2 | 715.7 |
| TMD2 | K1060T | 3179A>C | 383.2 | 614.0 | 1808.3 | 1128.6 | 2782.9 | 927.8 | 3104.3 |
| TMD2 | G1061R | 3181G>C | 52.6 | 65.0 | 249.2 | 201.8 | 546.2 | 62.1 | 365.4 |
| TMD2 | R1066C | 3196C>T | 45.7 | 48.3 | 205.0 | 194.1 | 363.4 | 84.6 | 340.9 |
| TMD2 | R1066H | 3197G>A | 56.4 | 81.3 | 298.1 | 207.0 | 834.2 | 68.0 | 549.4 |
| TMD2 | A1067T | 3199G>A | 281.5 | 493.7 | 1310.8 | 747.5 | 2405.3 | 636.4 | 3089.7 |
| TMD2 | G1069R | 3205G>A | 431.1 | 711.9 | 2196.8 | 1330.2 | 3182.2 | 1023.8 | 2888.1 |
| TMD2 | R1070W | 3208C>T | 185.2 | 284.8 | 662.3 | 430.7 | 1656.2 | 276.8 | 1986.6 |
| TMD2 | R1070Q | 3209G>A | 455.0 | 770.7 | 2375.7 | 1402.4 | 3362.0 | 1079.4 | 2995.7 |
| TMD2 | F1074L | 3222T>G | 171.5 | 366.4 | 792.9 | 371.2 | 1998.0 | 256.1 | 2094.1 |
| TMD2 | L1077P | 3230T>C | 53.9 | 63.8 | 243.3 | 205.5 | 556.9 | 69.2 | 415.3 |
| TMD2 | H1085P | 3254A>C | 49.1 | 51.7 | 212.5 | 201.6 | 422.9 | 52.1 | 352.6 |

| | | | | | | | | | |
|---|---|---|---|---|---|---|---|---|---|
| TMD2 | H1085R | 3254A>G | 63.6 | 86.7 | 291.6 | 214.0 | 803.4 | 63.1 | 701.9 |
| TMD2 | W1089X | 3266G>A | 48.8 | 57.2 | 241.3 | 205.9 | 395.7 | 70.2 | 347.5 |
| TMD2 | Y1092X | 3276C>A | 50.1 | 59.3 | 248.8 | 210.1 | 401.6 | 79.4 | 340.6 |
| TMD2 | Y1092X | 3276C>G | 49.4 | 58.3 | 248.4 | 210.3 | 417.7 | 67.5 | 323.0 |
| TMD2 | W1098R | 3292T>C | 54.0 | 63.3 | 246.3 | 210.2 | 404.0 | 63.8 | 307.5 |
| TMD2 | W1098C | 3294G>C | 55.4 | 76.0 | 287.1 | 209.3 | 817.8 | 61.1 | 624.1 |
| TMD2 | W1098C | 3294G>T | 64.0 | 94.7 | 327.2 | 221.3 | 818.4 | 56.4 | 572.6 |
| TMD2 | F1099L | 3297C>A | 113.3 | 305.3 | 776.0 | 288.0 | 2032.1 | 136.4 | 2158.9 |
| TMD2 | M1101K | 3302T>A | 51.9 | 62.4 | 242.6 | 201.6 | 405.8 | 62.5 | 368.7 |
| TMD2 | M1101R | 3302T>G | 47.0 | 54.3 | 257.7 | 223.1 | 412.4 | 67.8 | 365.0 |
| TMD2 | E1104X | 3310G>T | 47.5 | 53.6 | 219.4 | 194.7 | 389.8 | 59.6 | 318.5 |
| TMD2 | S1118F | 3353C>T | 214.9 | 371.1 | 860.8 | 498.6 | 2014.8 | 357.4 | 2446.1 |
| TMD2 | I1139V | 3415A>G | 411.6 | 701.1 | 2081.4 | 1221.8 | 3022.8 | 1045.1 | 3351.8 |
| TMD2 | D1152H | 3454G>C | 415.2 | 694.0 | 1966.9 | 1176.8 | 2860.6 | 1016.0 | 2914.4 |
| TMD2 | V1153E | 3458T>A | 148.4 | 194.8 | 396.8 | 302.3 | 979.7 | 148.5 | 972.9 |
| TMD2 | L1156= | 3466T>C | 418.4 | 781.0 | 2424.3 | 1299.0 | 3166.7 | 1011.0 | 2753.5 |
| TMD2 | L1156= | 3468G>A | 452.7 | 748.5 | 2163.2 | 1308.2 | 3213.4 | 1114.5 | 2955.7 |
| TMD2 | L1156F | 3468G>T | 429.7 | 765.3 | 2222.8 | 1248.2 | 3263.2 | 1046.3 | 3034.5 |
| TMD2 | L1156= | 3468G>A | 416.1 | 695.5 | 2192.4 | 1311.9 | 3186.0 | 1050.1 | 3414.1 |
| TMD2 | R1158X | 3472C>T | 56.5 | 72.4 | 281.5 | 219.8 | 447.2 | 79.9 | 345.4 |
| TMD2 | S1159F | 3476C>T | 305.9 | 551.5 | 1424.1 | 789.8 | 2380.9 | 689.4 | 2563.5 |
| TMD2 | S1159P | 3475T>C | 393.1 | 761.1 | 2114.6 | 1092.2 | 3265.2 | 930.5 | 3404.4 |
| NBD2 | R1162X | 3484C>T | 51.0 | 56.9 | 230.0 | 206.1 | 402.3 | 80.3 | 308.1 |
| NBD2 | R1162L | 3485G>T | 309.0 | 496.2 | 1330.1 | 828.4 | 2337.6 | 705.6 | 2391.8 |
| NBD2 | S1196X | 3587C>G | 318.2 | 437.3 | 1322.6 | 962.4 | 2054.8 | 764.6 | 1819.5 |
| NBD2 | W1204X | 3611G>A | 327.5 | 413.6 | 1226.5 | 971.1 | 1896.7 | 838.2 | 1871.5 |
| NBD2 | W1204X | 3612G>A | 294.3 | 338.0 | 1052.3 | 916.3 | 1736.5 | 654.1 | 1301.8 |
| NBD2 | I1234V | 3700A>G | 420.4 | 727.9 | 2304.5 | 1331.1 | 3352.8 | 1119.6 | 3264.0 |
| NBD2 | S1235R | 3705T>G | 427.2 | 773.8 | 2436.0 | 1344.9 | 3377.5 | 1069.0 | 3248.1 |
| NBD2 | V1240G | 3719T>G | 259.0 | 561.6 | 1454.7 | 671.0 | 1793.5 | 759.4 | 2301.6 |
| NBD2 | G1244E | 3731G>A | 445.1 | 751.4 | 2281.9 | 1351.9 | 3353.1 | 1078.5 | 3027.0 |
| NBD2 | T1246I | 3737C>T | 439.7 | 741.3 | 2179.2 | 1292.6 | 3285.8 | 1041.2 | 2973.8 |
| NBD2 | G1249R | 3745G>A | 414.4 | 730.8 | 2150.7 | 1219.5 | 3086.7 | 1049.5 | 3486.7 |
| NBD2 | S1251N | 3752G>A | 424.3 | 833.9 | 2660.0 | 1353.3 | 3312.6 | 1103.4 | 2909.7 |
| NBD2 | S1255P | 3763T>C | 422.0 | 712.3 | 2123.7 | 1258.2 | 3161.5 | 1066.1 | 3152.1 |
| NBD2 | I1269N | 3806T>A | 225.1 | 325.3 | 800.2 | 553.7 | 1418.7 | 506.9 | 1529.0 |
| NBD2 | D1270N | 3808G>A | 412.4 | 699.6 | 2106.7 | 1241.8 | 3151.4 | 998.4 | 2873.2 |
| NBD2 | W1282X | 3846G>A | 114.0 | 150.2 | 464.8 | 352.7 | 899.7 | 174.9 | 712.3 |
| NBD2 | W1282R | 3844T>C | 145.0 | 170.8 | 428.1 | 363.3 | 949.0 | 225.5 | 916.7 |
| NBD2 | R1283M | 3848G>T | 188.2 | 234.2 | 525.8 | 422.4 | 1150.6 | 362.9 | 1436.7 |
| NBD2 | R1283S | 3849G>C | 192.0 | 255.2 | 617.6 | 464.6 | 1219.6 | 343.1 | 1476.5 |
| NBD2 | Q1291H | 3873G>C | 438.7 | 735.9 | 2147.7 | 1280.1 | 3235.2 | 1075.4 | 2985.2 |
| NBD2 | Q1291R | 3872A>G | 438.0 | 777.1 | 2464.0 | 1388.8 | 3453.4 | 1154.4 | 3497.3 |
| NBD2 | V1293G | 3878T>G | 431.3 | 720.1 | 2223.0 | 1331.4 | 3314.1 | 1118.6 | 3486.8 |
| NBD2 | N1303K | 3909C>G | 88.4 | 91.7 | 271.8 | 261.8 | 609.5 | 118.4 | 506.6 |
| NBD2 | Q1313X | 3937C>T | 50.3 | 57.5 | 231.0 | 202.2 | 403.6 | 72.6 | 335.0 |

| | | | | | | | | | |
|---|---|---|---|---|---|---|---|---|---|
| NBD2 | L1324P | 3971T>C | 80.7 | 78.4 | 241.7 | 248.6 | 548.5 | 106.9 | 536.9 |
| NBD2 | L1335P | 4004T>C | 84.5 | 79.9 | 229.6 | 242.9 | 563.9 | 100.9 | 446.9 |
| NBD2 | G1349D | 4046G>A | 416.5 | 682.8 | 2008.6 | 1225.3 | 3062.5 | 1013.7 | 2839.5 |
| NBD2 | I1366N | 4097T>A | 168.8 | 206.7 | 499.7 | 408.1 | 1048.4 | 249.7 | 993.5 |
| NBD2 | H1375P | 4124A>C | 405.3 | 657.0 | 2030.5 | 1252.7 | 3171.4 | 968.5 | 3070.1 |
| NBD2 | L1480P | 4439T>C | 383.9 | 621.1 | 1747.4 | 1080.3 | 2658.3 | 964.3 | 3053.1 |
